# Tackling Bias in Cortical Thickness Estimation in UK Biobank Using Harmonisation Approaches

**DOI:** 10.64898/2026.05.22.726536

**Authors:** Jacob Turnbull, Gaurav Bhalerao, Rach Dawson, Frederik Lange, Fidel Alfaro-Almagro, Stephen Smith, Ludovica Griffanti

## Abstract

Big neuroimaging data enable researchers to study subtle structural and functional brain changes and relationships between brain characteristics and genetics, lifestyle, and disease factors. However, substantial effort is needed to minimise technical, non-biological differences between data batches to avoid incorrect inferences. In this study, we address a previously identified bias in UK Biobank FreeSurfer IDPs derived from only the T1 image compared to those using both T1 and T2-FLAIR by treating the bias as a batch effect and using harmonisation approaches. We investigate and characterise this bias through direct within-participant comparison at the image and IDP level, comparing the results with those seen in the wider UKB sample. We then assess different methods of addressing the effect of missing T2-FLAIR, starting from simple linear regression before moving to ComBat, a widely used harmonisation method, testing different approaches for applying ComBat and showing its similarity to simple linear regression. Finally, we examine how ComBat estimates vary with batch and sample size. Our results show clear benefits in using both T1 and T2-FLAIR data in FreeSurfer, as opposed to just the T1, which is more common, with the pial surface fitting being less likely to fail and showing greater biologically plausible inter-subject variability. This is particularly important for cortical thickness IDPs, where T2-FLAIR omission leads to reduced true variability and systematic underestimation, as shown through within-participant repeat testing. We demonstrate that ComBat can address this bias, with its standard use (i.e., applied separately on different IDP categories) showing the best improvement in cortical thickness measures where the bias is strongest, and we find that it is important not to pool ComBat priors across different classes of IDPs. Our proposed version of ComBat with a reference batch (i.e., estimating mean and variance only from data with T2-FLAIR available) performed best in recovering both mean and variance differences between batches across different IDP classes and offers a promising approach for cases where a reference batch is clearly identifiable. While ComBat reliably corrects mean (additive) batch effects with relatively small sample sizes (*≈*30 subjects per batch), we show that its variance (multiplicative) correction is substantially less stable, requiring much larger sample sizes and becoming unreliable when batches are small or imbalanced, or when there is a large variance difference between them.

## 1 Introduction

Over the past decade, there has been a growing push towards increased data sharing and collaboration within the neuroimaging research community. This increased data sharing and collaboration, along with improvements in the resolution, availability of modalities and overall quality of neuroimaging datasets, has resulted in the production of large and diverse datasets, often referred to as “big data”. These large datasets enable the study of anatomical and functional changes of the brain and their potential relationships with genetic and lifestyle factors, as well as the identification of early markers and potential mechanisms of disease, which is only possible through such a large number of samples (S. M. Smith & Nichols, 2018).

Broadly, such large neuroimaging datasets can be created in one of three ways. The first is through the accumulation of a large number of scans with the same protocol and scanner. For example, the Human Connectome Project (HCP), which collected extensive multimodal MRI data from over 1,000 young healthy volunteers (Van Essen et al., 2012), or UK Biobank (UKB), a large-scale epidemiological study which has recently completed the multimodal MRI data collection of 100,000 members of an original cohort of 500,000 (Miller et al., 2016; Sudlow et al., 2015). While single-site accumulation studies have relatively low technical variability, they are incredibly expensive, time-consuming, and often use population samples that are not representative of the general population. Additionally, unforeseen part replacements or scanner breakdowns can cause bias that can be hard to address.

The second is through multisite prospective data collection, which uses the same imaging protocol at multiple sites and scanners. This method is often more scalable than the first approach, and the homogeneous protocol helps to focus data differences on biological variability. Some examples include the Alzheimer’s Disease Neuroimaging Initiative (ADNI-1, ADNI-2) (Petersen et al., 2010) and the Ageing Brain Cohort (ABC) (Newman-Norlund et al., 2021). However, even with standard protocols and acquisitions, differences in scanner hardware, software and pulse sequences still need to be addressed.

The third method is through retrospective data merging, where datasets that were collected on different scanners, with often very different imaging protocols, are combined to make use of the wealth of data available in the field, which is only possible due to data sharing and international collaboration becoming more common. Two major examples are the ENIGMA consortium (Enhancing NeuroImaging Genetics through Meta Analysis) (Thompson et al., 2020), which has pooled data from over 50 working groups across the world with a large range of neurological conditions, and the iSTAGING consortium, which has aggregated the data of participants with mild cognitive impairment from multiple large neuroimaging studies (Habes et al., 2021). This retrospective data-merging approach typically has the most severe technical variability, often making it difficult to pool these datasets together due to inconsistent resolution and image quality protocols (Cetin Karayumak et al., 2019).

MRI harmonisation is the general term given to any method of reducing technical, generally non-biological differences between batches of MRI data, where batches can represent any grouping variable, such as scanner or site. Strategies for harmonising MRI data can be applied prospectively during the imaging stage through the use of the same scanner and/or acquisition protocol and at the data processing stage through the use of standard image processing pipelines, as well as through retrospective harmonisation algorithms, which aim to remove bias in the extracted MRI measures arising from technical variability while preserving biological signal. When discussing harmonisation approaches throughout this paper, we are only considering retrospective harmonisation algorithms. The differences between datasets arising from non-biological variability are often referred to as “batch effects”, with batch being any grouping variable that can be used to describe differences in the acquisition or processing of the different measurements.

UK Biobank is the world’s largest neuroimaging study, having recently completed the initial 100,000 multimodal brain scans and now continuing with the collection of 60,000 repeat scans. The UKB dataset provides a unique opportunity to study the relationship between detailed, multimodal neuroimaging data of the brain and a wealth of genetic, lifestyle, health and cognitive measures, as well as to study human brain ageing. MRI data is collected on identical scanners and processed using an automated image processing pipeline (Alfaro-Almagro, 2021) which removes artefacts and performs quality control checks on each individual’s datasets, making them comparable between participants across modalities, before extracting a large quantity of imaging-derived phenotypes (IDPs) describing observable structural and functional characteristics of the brain (Elliott et al., 2018).

However, even with a study which is as standardised as UKB, technical variability can still occur and needs to be carefully addressed before downstream analysis. While there is substantial effort to ensure all participants in UKB have the same modalities available, there are sometimes interruptions or issues that cannot be controlled for during acquisition, resulting in missing modalities in some cases. As a downstream effect, this might mean that a particular IDP cannot be derived, or that the calculation by which the IDP is estimated may follow a different pathway which may introduce a batch effect.

FreeSurfer analysis of brain structure is, where possible, performed using both T1 and T2-FLAIR images, obtaining complementary value from those two structural modalities. However, if the T2-FLAIR is not available, FreeSurfer is run using just the T1 (Fischl, 2012; Iglesias et al., 2018). In previous works, it has been shown that when the T2-FLAIR image is absent in FreeSurfer processing, there is a higher rate of surface fitting failure and potential for bias in some of the resulting IDPs (Ferreira-Atuesta et al., 2022; Lindroth et al., 2019).

In a broad analysis of confounding effects in UKB, Alfaro-Almagro et al. (2018) showed that when linearly modelled with a larger set of explanatory variables, missing T2-FLAIR in FreeSurfer processing, henceforth referred to as the FST2 effect, accounted for up to 14% of the variance explained and 7% of unique variance explained in some structural FreeSurfer IDPs. In the most affected IDPs, the variance explained by the FST2 effect was higher than by well-established variables such as age or sex, despite the portion of participants having FreeSurfer run with only T1 being relatively low (<3% of the total sample). Depending on the approach, deconfounding and harmonisation share several similarities but differ primarily in their scope and intended use. Deconfounding methods, such as regression, are typically applied to derived measures individually to remove the effects of variables of non-interest. In contrast, harmonisation methods operate across multiple features simultaneously, aligning batch-specific distributions to a common latent space across the dataset, effectively changing the data to improve comparability. In essence, deconfounding is typically limited to being subtractive, whereas harmonisation may use a more general transformation.

In this work, we treat the absence of T2-FLAIR in FreeSurfer processing as a batch effect (Bayer et al., 2022). Here, we adopt a broad definition of batch effects as any systematic, non-biological differences arising from acquisition, processing, or analysis pipelines. While batch effects are often associated with site or scanner differences, they can equally arise from differences in processing inputs or configurations. In this case, the omission of T2-FLAIR represents a consistent and well-defined processing difference affecting a substantial subset of participants (>1000), making it appropriate to model and correct using harmonisation approaches rather than treating it as random missingness or excluding affected subjects.

In this study, we aim to achieve several objectives. The first is the systematic evaluation of the effect of not including T2-FLAIR in FreeSurfer processing. While it has been previously noted that T2-FLAIR’s inclusion can be beneficial for avoiding failures when fitting the pial surface and for preserving age-related atrophy trends in cortical thickness measures (Ferreira-Atuesta et al., 2022; Lindroth et al., 2019), the overall difference between IDPs has not been systematically evaluated, nor has the impact on downstream associations between IDPs and non-imaging variables. Here we aim to demonstrate how including T2-FLAIR in FreeSurfer can be beneficial for more accurate and representative IDPs and to show how not including it can potentially bias downstream analysis.

The second is the production of a batch-corrected set of IDPs through the modelling and removal of the FST2 effect from UKB FreeSurfer IDPs. In previous studies, these participants have often been excluded from analysis; by correctly removing the bias from missing T2-FLAIR, we generate a set of batch-corrected structural IDPs and subsequent analysis with non-imaging variables that include them. We also aim to show the differences between ComBat harmonisation and standard deconfounding using regression. We assess each difference between ComBat and simple regression directly, showing the statistical impact of each additional ComBat term, before comparing different ways ComBat can be applied (Fortin et al., 2018; Marzi et al., 2024). We aim to identify the best way to produce these corrected IDPs and to inform its implementation in future core processing pipelines.

Finally, we seek to utilise the large sample size in UKB to examine how variable the batch corrections in ComBat are at different sample sizes and batch sizes. Previous studies have examined how well harmonisation is achieved at different sample sizes through the downstream statistical analysis of the batch effect, but here we aim to explore how well estimated the ComBat corrections are and if they can generalise between different sample sizes (Orlhac et al., 2022; Parekh et al., 2022). In doing so, we provide an assessment of how suitable ComBat is for extending to smaller datasets, providing a modified strategy that uses a reference site as well as assessing whether it is appropriate for research scenarios using machine or deep learning, where test and train splits are essential to avoid data leakage and overfitting.

## 2 Methods

### 2.1 Ethics

This research was conducted using data from UK Biobank under approved application number 8107. UK Biobank has ethical approval from the Northwest Multicentre Research Ethics Committee as a research tissue bank, and all participants provided informed consent for their de-identified data to be used for health-related research in the public interest. This study involved secondary analysis of de-identified UK Biobank data and was conducted in accordance with the UK Biobank Access Procedures.

### 2.2 Dataset

The total number of datasets available was 72,288. Of these participants, we initially included all those with a usable FreeSurfer output. This initial filtering resulted in the exclusion of 2,178 participants who had no available FreeSurfer output, with this exclusion largely being due to being flagged by QoalaQT (a tool to perform QC on FreeSurfer outputs) and then manually checked, or the run time of FreeSurfer being too long for these participants (i.e., not converging) (Alfaro-Almagro, 2021; Klapwijk et al., 2019). Of the remaining 70,110 participants, 68,646 had FreeSurfer run with both T1-weighted and T2-FLAIR images, and 1,464 participants had it run with only T1. Of the participants who had FreeSurfer run without the T2-FLAIR image, the majority of omissions were caused by missing or low-quality T2-FLAIR scans. 464 participants had no available T2-FLAIR scans, with only 23 of these participants having an available diffusion MRI scan, implying that in the majority of these cases, acquisition was interrupted. An additional 482 T2-FLAIR scans were deemed unusable by quality control checks run before FreeSurfer, and 543 cases had an early version of the UKB acquisition protocol, which is incompatible with the current image processing pipeline. Finally, one participant had incorrect image dimensions in their T2-FLAIR image, and an additional 4 had only T1 used for unspecified reasons. Some of these datasets may have had more than one of the above issues mentioned. Additionally, as some data was from repeat scans of participants already present, we removed these measures and took only the most recent data, leaving 62,823 unique participants with 1,409 missing T2-FLAIR.

The IDPs used were all FreeSurfer IDPs, totalling 1,272 unique measures of five different major categories: regional and tissue volume (483), cortical area (372), cortical thickness (306), regional and tissue intensity (41), and cortical grey-white matter contrast (70). Four IDPs corresponding to the volume and intensity of the septum pellucidum, referred to as the 5th ventricle in FreeSurfer, and to the volume and intensity of non-white matter hypo-intensities had missing or zero data in over 80% of participants. These four IDPs were subsequently excluded, as the vast majority of participants would have measures that are too small or did not have these structures; as such, their inclusion would cause biased estimates in the variance of the data (Fischl et al., 2002). A more extensive overview of the atlases and tools used by FreeSurfer within the UKB pipeline can be found on the FreeSurfer wiki for version 6.0 (https://surfer.nmr.mgh.harvard.edu/fswiki) and described in section 3.6 of the UKB brain imaging documentation (https://www.fmrib.ox.ac.uk/ukbiobank/).

The IDPs were then converted to median-centred Z-scores, and all participants with a score of Z > |8| in any IDP were considered extreme outliers and omitted from the dataset to avoid large bias. This totalled a removal of 18,305 participants who had a Z-score greater than |8| in one or more structural IDPs, leaving the final number of participants in this study as 44,535, with 983 of these participants missing the T2-FLAIR image. This conservative approach of excluding entire subjects was used to maximise the interpretability of the results. In general, it is more common to exclude individual outlier IDPs.

### 2.3 Characterising the batch effect

In order to assess directly the impact of the FST2 batch effect, we began by taking 100 randomly drawn participants from the group with a T2-FLAIR image and re-running the FreeSurfer part of the UKB image processing pipeline on this subgroup using only their T1-weighted image. By rerunning FreeSurfer without T2-FLAIR, we created a paired participant group with which we can directly assess the individual impact of not including T2-FLAIR in FreeSurfer processing. We qualitatively compared the pial surface and grey-white matter boundaries of the original outputs (the outputs when both T1 and T2-FLAIR are used) against the new outputs, visually inspecting the difference in the surfaces when overlaid on each image modality.

We then extracted IDPs from the re-run FreeSurfer outputs and directly compared them against the original outputs. We first looked at the specific effect on the mean (often called a location shift) by calculating Cohen’s d between the two datasets, as seen in equation (1). Here we define any value above 0.6 as a large effect size, with a measure between 0.2 and 0.6 being classified as a medium effect size, and anything less than 0.2 being a small effect size, as consistent with the widely accepted values in the literature (Riffenburgh & Gillen, 2020b):

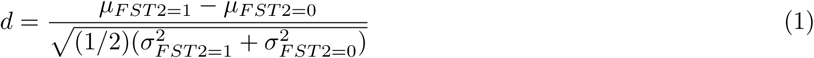

Next we looked at scaling differences between the two groups for each IDP. To do this, we took the ratio of the two batches’ variances for each IDP as described in equation (2). Here, a *V_r_* equal to 1 would indicate identical variance between batches.

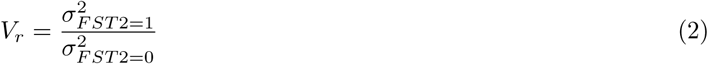

To quantify the impact of processing with versus without T2-FLAIR, we examined subject-specific paired differences in cortical thickness estimates. For each subject and region, we computed the absolute difference (in mm) between the original IDPs and those derived from the pipeline excluding T2-FLAIR, providing a direct measure of the magnitude of processing-induced variation. We then calculated the correlations of FreeSurfer IDPs with each other, with the hypothesis that better correlations between IDPs may indicate improved consistency of processing (Lindroth et al., 2019). This was done independently for different types of IDPs (i.e., correlations between thickness IDPs rather than thickness IDPs with volume IDPs).

After looking at these metrics on the paired participant group, we applied the same tests on the wider UKB sample (i.e., the unpaired data). This was done to verify that the observed bias effects on the mean and scaling were replicated in the unpaired sample.

### 2.4 Harmonisation and deconfounding approaches

After quantifying the effect on the mean and the variance due to the FST2 batch effect, we applied different decon-founding and harmonisation approaches to address it. In this work, we distinguish between deconfounding (removal of covariate effects via regression) and harmonisation (alignment of batch distributions, including both mean and variance adjustments), noting that some methods (e.g., ComBat) perform both simultaneously. We start with the simplest application of linear regression for deconfounding and add terms to the model in a stepwise manner so each subsequent approach has a clear comparison to the previous ones. We do this to show the direct differences between each component in ComBat compared to simple deconfounding before showing different ways in which one might apply ComBat for harmonisation.

#### 2.4.1 Method 1: Linear Regression of standard confounds

Before directly addressing the FST2 effect, we first used simple linear regression to perform deconfounding of some other variables. We chose age, sex, head position (in the scanner) and head size as the four confounds to be modelled, as they were found by (Alfaro-Almagro et al., 2021) to explain a large portion of unique “uninteresting” variance within structural IDPs. By first modelling the data without including the binary FST2 indicator, we establish that observed statistical batch effects aren’t caused by different underlying distributions (for example, age and sex differences correlating with batch), as well as establishing a baseline approach against which to compare different harmonisation methods that do not directly remove the FST2 effect.

The regression model we use is shown in equation (3). Here **X_i_** is the design matrix containing the subject-specific measure for each of the confounds and *β***_j_** are the beta coefficients from ordinary least squares (OLS), with *i* denoting the participant and *j* denoting the IDP index. The beta coefficients were estimated using the pseudoinverse, *β_j_* = (*X^T^ X*)^−1^*X^T^ y_j_*, as used in the ComBat algorithm. The terms *α_j_* and *ɛ_ij_* represent the feature-specific intercept term not included in *X_i_* and the subject-specific noise for IDP *j*, respectively (see Section 2.3.3).

Here, we define the “standard” set of confounds as age, sex, head size scaling, and head position in the scanner’s Z-axis (the longitudinal field direction), with all variables being mean-centred.

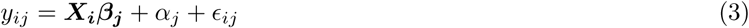

#### 2.4.2 Method 2: Linear regression including batch

We then extended method 1 to include the missing T2-FLAIR (FST2) as a binary indicator of batch in the design matrix. In doing so, we estimate beta coefficients for each IDP and residualise the data while accounting for batch effects, with the change being shown in equation (4). When calculating the beta coefficients for the FST2 variable, we include it in the main **X_i_** matrix but show it separately in equation (4) for readability.

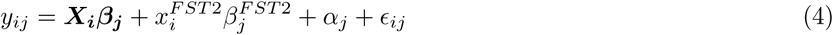

#### 2.4.3 ComBat

ComBat (Combating Batch effects) is an empirical Bayes method for harmonising multi-site/batch data that was originally introduced by Johnson et al. (2007), for correcting batch effects in microarray gene expression data before being adapted for uses in medical imaging in Fortin et al. (2017, 2018). Since then, it has been widely adopted as a “silver standard” of image harmonisation, and a large number of ComBat variations have been created, such as long-ComBat for longitudinal data, ComBat-GAMs for non-linear covariate effects, and auto-ComBat, which uses image-level metadata to automatically determine batch labels (Carré et al., 2022; Orlhac et al., 2022; Pomponio et al., 2020).

Since part of our goal for this study was to provide a systematic evaluation of each of the ComBat parameters and provide a direct comparison with linear regression, we used the most frequently used version of ComBat for MATLAB available (Fortin (2021)). The use of this openly available version facilitates reproducibility as well as the implementation of the variations described below, which we have made available in both Python and MATLAB in the supplementary materials.

ComBat models the data as defined in equation (5), with the additional index *g* denoting the batch to which the participant *i* belongs. Here *y_ijg_* is the observed measure for participant *i*, feature *j* and batch *g*, *X_i_β_j_* is the standard set of confounds design matrix and associated beta coefficients from ordinary least squares; *γ_jg_* is the feature- and batch-specific additive term; and *δ_jg_* is the feature- and batch-specific term on the participant-specific signal *ɛ_ij_*. A key limitation of this formulation is that *ɛ_ij_* conflates participant-specific biological signals with residual noise. Because both components are scaled jointly by *δ_jg_*, the model does not explicitly distinguish between true subject-level variation and stochastic measurement error. As a result, the batch-specific scaling term implicitly acts on both signal and noise simultaneously, rather than selectively correcting only unwanted variation. This reflects the underlying assumption of ComBat that the residual variation follows a common distribution across batches, meaning that participant-specific effects are not separately identifiable from noise within this framework.

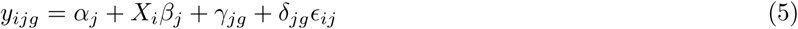

The ComBat algorithm first standardises the data by subtracting the estimated feature-wise means and covariate effects before dividing by the feature-wise variance, such that the data can be assumed to have the form shown in equation (6).

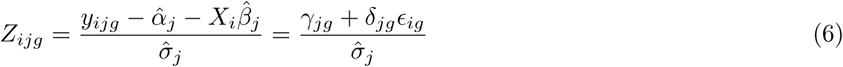

#### 2.4.4 Method 3: ComBat with no variance correction

We first applied a reduced form of ComBat in which the multiplicative (variance) adjustment term is omitted, such that only the additive batch effect is modelled. In contrast to a pooled implementation across all IDPs (Method 4), this method was applied separately within each IDP class (e.g., cortical thickness, volume, and area), consistent with the standard use of ComBat in neuroimaging studies, where features of the same type are harmonised together. The estimate of 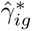 depends on 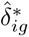 and vice versa. Consequently, these parameters are obtained iteratively, as no closed-form solution exists. Estimation proceeds sequentially, beginning with an initial posterior estimate for 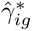 (obtained via the pseudoinverse of the batch design matrix applied to the standardised data), which is then used to update 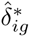. This process is repeated until convergence is achieved. The harmonised data are defined as seen in equation 7, where *k* indexes IDP class.

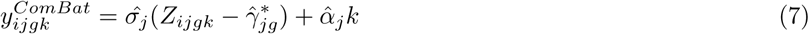

Although ComBat’s iterative Bayesian procedure still uses the estimate for the scaling term to inform the value for the location (mean shift) parameter, it has no direct impact on the final adjustment in this formulation (and hence is not identical to simple batch confound regression). This formulation allows us to isolate the impact of the empirical Bayes estimation of the additive batch effect while retaining class-specific priors and hyperparameters. By constraining estimation within IDP classes, the prior distributions are expected to more closely reflect the underlying structure and scale of the data, potentially improving robustness when batch effects differ across feature types (e.g., thickness vs intensity contrast IDPs). We consider this variant first because it is conceptually closest to regression-based deconfounding, allowing us to isolate the effect of the estimated location term, rather than relying on regression coefficients. A full derivation of the ComBat model is provided in Johnson et al. (2007).

#### 2.4.5 Method 4: ComBat with scaling correction, applied to all IDPs at once

In contrast to the standard application of ComBat, which is typically performed on a single class of features (e.g., cortical thickness or volumetric measures), we also implemented a pooled version of ComBat, in which all IDPs were harmonised jointly under a single model (hence mimicking a naive default application of ComBat to a large set of imaging features spanning multiple feature types). In this formulation (equation 8), ComBat is applied across all features simultaneously, such that the empirical Bayes priors governing the batch-specific parameters are estimated using the full set of IDPs:

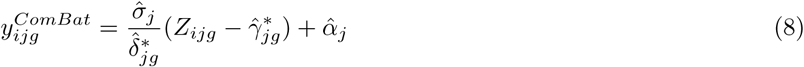

Here, all features *j* are treated as arising from a shared prior distribution over the batch-specific parameters *γ_jg_* and *δ_jg_*. This pooled implementation is not standard practice in neuroimaging harmonisation, where ComBat is usually applied within homogeneous feature sets. However, we include it here for two primary reasons. First, the source of the batch effect considered in this study—the omission of T2-FLAIR during FreeSurfer processing—represents a shared technical bias affecting all IDPs, arising from differences in surface fitting. This provides a methodological justification for considering whether a shared prior across IDPs may capture this global source of variation. Second, pooling IDPs allows us to directly examine the effect of the empirical Bayes prior structure on the estimated batch corrections. When ComBat is applied within IDP classes, each class has its own prior distribution and associated hyperparameters, which are estimated from a relatively homogeneous set of features. This is expected to result in tighter priors and stronger shrinkage of the batch-specific parameters. In contrast, when applied across heterogeneous IDPs, the prior distribution may become substantially broader, reflecting differences in scale and variability between feature types. In this case, the empirical Bayes shrinkage may be weaker, and the resulting corrections may more closely resemble feature-wise estimates.

Comparing this pooled implementation with the standard class-wise approach therefore allows us to assess how the structure and specificity of the prior influence the stability and magnitude of the ComBat corrections. While previous work has suggested that pooling features across classes may have limited impact on downstream analyses (Radua et al., 2020), we explicitly evaluate this assumption in the context of a known, technically-driven batch effect.

#### 2.4.6 Method 5: ComBat applied separately on different IDP categories

We then applied ComBat separately within distinct classes of IDPs (cortical thickness, cortical area, cortical volume, intensity and grey-white matter contrast). As in equation 7, the additional index *k* is used to denote the IDP class to which ComBat is applied.

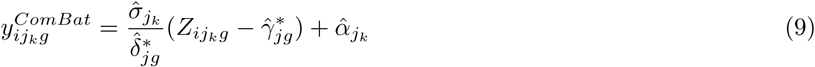

This class-wise implementation reflects the standard use of ComBat in neuroimaging, where harmonisation is typically performed on a single measurement type, with the only difference being the removal rather than the preservation of *X_i_β_j_*. By estimating the empirical Bayes priors within each IDP class, the model leverages information from a relatively homogeneous set of features, which is expected to produce more informative and tighter prior distributions for the batch-specific parameters. This approach may be advantageous when different classes of IDPs exhibit distinct distributions, scales, or sensitivities to batch effects, as the class-specific priors can better capture these differences. In particular, for features strongly affected by the batch effect (e.g., cortical thickness), this may result in more stable and appropriate shrinkage of the estimated batch corrections.

#### 2.4.7 Method 6: ComBat with a reference batch

Our final implementation uses a “reference-batch” version of ComBat (M-ComBat). Rather than standardising using global estimates across all batches, we standardise the data using the mean and variance of a single reference batch. This places all batches on the reference batch scale before estimating the ComBat location and scale adjustments, *γ*^∗^ and *δ*^∗^. After these corrections are applied, the data are rescaled using the same reference batch mean and variance. We apply ComBat separately within each IDP category so that the empirical Bayes priors and hyperparameters are estimated within feature-homogeneous groups, as done with methods 3 and 5.

This differs from the formulation proposed by Stein et al. (2015), which initially standardises each batch using its own mean and variance before rescaling the adjusted data to a chosen gold-standard batch. In our implementation, the reference batch defines the scale throughout, so the harmonised data are directly anchored to that batch.

Conceptually, this is the same as anchoring one batch to another so that the anchor batch is unchanged during harmonisation, and the other batches are pulled towards the reference batch’s mean and variance. This modified ComBat approach requires that one batch be treated as a “ground truth” to which all other batches should be harmonised. We set the batch with T1 and T2-FLAIR in FreeSurfer as our reference batch, as it is substantially larger and generally considered to be more accurate. This assumption is supported by the observed reduction in surface failures and higher between-subject variability. In this case we hypothesise that M-ComBat may offer some advantages over regular ComBat. In some situations identifying a reference batch is not possible or appropriate, for example, where there are many batches of similar sample sizes. The initial data standardisation step and final adjustment are shown below in equations 10 and 11, respectively.

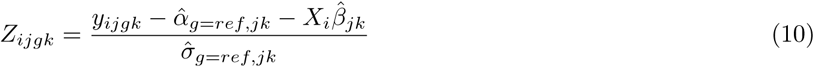

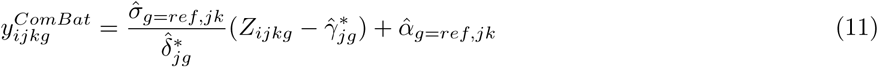

The reference batch will have a scaling term 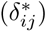 equal to 1 and a location term 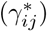 equal to zero, making the final adjustments of the reference site equivalent to method 2, omitting the FST2 term.

To summarise, the six methods of handling confounds/harmonising compared in this work are:

- Method 1: Linear regression of the effects associated with the standard set of confounds only.
- Method 2: Linear regression of the effects associated with the standard set of confounds and an additional binary vector indicating FST2 batch.
- Method 3: ComBat handling of both the standard confounds and the FST2 batch effect with no scaling term in the model, applied separately within IDP classes.
- Method 4: ComBat handling of both the standard confounds and the FST2 batch effect, applied to all IDPs at once. Method 4 represents a non-standard implementation used to assess the effect of pooling IDPs across classes under a shared prior.
- Method 5: ComBat handling of both the standard confounds and the FST2 batch effect applied separately within IDP classes.
- Method 6: ComBat handling of confounds and the batch effect, setting the batch with T2-FLAIR in FreeSurfer (FST2=1) as the ground-truth reference batch to be harmonised to, applied separately within IDP classes.

### 2.5 Harmonisation efficacy

MRI harmonisation is considered successful when it fulfils two criteria. The first is the removal of the batch effect so that previously uncalibrated measures can be directly compared. The second is the preservation of important biological information, often shown using the preservation of the relationship between the data and simple variables like age or sex (Dewey et al., 2022; Lu et al., 2025).

#### 2.5.1 Batch effect removal

To show the batch effect and test its subsequent removal in each method, we use Cohen’s d and variance ratio (*V_r_*) as defined in equations 1 and 2. In doing so, we remain consistent with our previous method of characterising the batch effect in both the repeated group and the larger UKB sample, as well as being consistent with the ComBat model (which applies a batch correction on the mean and spread of the data).

Additionally, as the goal of harmonisation is to make different batches more comparable to each other, we include a two-sample Kolmogorov-Smirnov (KS) test on each IDP (Riffenburgh & Gillen, 2020a). The two-sample KS test calculates the difference in the empirical distribution of each sample using a supremum function, which compares directly the difference of a function at each point and takes the maximum value, as shown in equation (12). This test does not assume a specific distribution of the data and is sensitive to differences in both the mean and scale between batches.

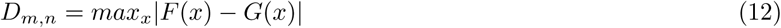

Here, m and n represent the sample size of the FST2=1 and FST2=0 batches, respectively, with F(x) and G(x) being their observed cumulative distributions. The null hypothesis assumes that both batches come from the same population and is rejected if the distance is larger than the critical value shown in equation (13), at a p-value of *α*. Here, a rejection of the null hypothesis is indicative of a statistically significant difference in the distributions of the two batches.

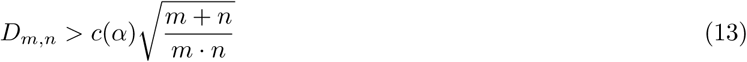

Here, *c*(*α*) is the inverse of the KS distribution, and it acts as a scaling factor on the required difference (*D_m,n_*). The value of *c*(*α*) at a given alpha is obtained from the Kolmogorov distribution (tabulated or approximated analytically by 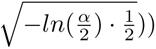. As such, the final condition for the rejection of the null hypothesis at a significance level of *α* is shown in equation (14).

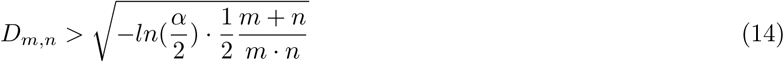

#### 2.5.2 Ensuring preservation of biological information

Typically, the preservation of biological information is shown by looking directly at the relationship between the measures and biological covariates, such as age or sex, before and after harmonisation. This can be done through the correlations with measures being unchanged or improved after harmonisation, improved prediction accuracy of a machine learning classifier, or through some other analogous method (Marzi et al., 2024).

While many harmonisation studies assess biological signal preservation using known associations, for example with age or sex, these effects are both strong and well-characterised in UK Biobank and are not expected to be badly confounded by the FST2 batch effect. As such, their preservation does not provide a sufficiently sensitive test of harmonisation performance in this context. Instead, we focus on the preservation of more subtle associations between IDPs and a large set of non-imaging variables (nIDPs), which better represent the analytical value of large-scale datasets such as UK Biobank. By examining whether these weaker, higher-dimensional relationships are maintained after harmonisation, we provide a more stringent and practically relevant validation of biological signal preservation. Previous works have shown that modelling and removing confounding effects can reveal and strengthen subtle but statistically significant relationships between IDPs and nIDPs (Alfaro-Almagro et al., 2021). This was part of the motivation in the deconfounding of age, sex, head position, and head size scaling performed by each method, as it allows us to ensure that these subtle relationships aren’t removed during harmonisation and also to identify potential spurious correlations that may be driven by the FST2 effect. Overall, we expect that the vast majority of correlations will be preserved between our methods, with this being especially important for the strongest correlations between IDPs and nIDPs. By showing that these effects are preserved, we can infer that true biological signals of importance are preserved in the data.

We looked at the associations between the IDPs after applying each of our methods and a set of over 17,000 nIDPs, with the effects from the standard set of confounds being modelled out, using Pearson’s correlation coefficient. The correlation coefficients were then transformed using the Fisher R-to-Z transform in order to stabilise variance and approximate normality, and the results were expressed as Bland-Altman plots of agreement (Corey et al., 1998; Giavarina, 2015).

We compare the 6 methods against each other, expressing the results as Bland-Altman plots of agreement over all IDP-nIDP pairs. We compared methods 1, 2 and 3 against each other to show directly the change in correlations when removing the mean shift component of the FST2 effect and to see whether the empirical Bayes handling of the effect on the mean performed by ComBat showed any major differences when compared with linear regression. We then compared methods 3 and 4 to assess the impact of the scaling term in ComBat before finally comparing method 4 to methods 5 and 6. This approach allows us to assess whether applying ComBat to distinct IDP categories or applying it with a reference site has any meaningful impact on the correlations with nIDPs.

### 2.6 Investigating generalisability of ComBat across batch and sample size

ComBat harmonisation consists of two primary steps. First, biological and technical covariate effects are estimated and removed via regression, after which the residualised data are standardised using estimates of the global (non–batch-specific) mean and variance for each feature. Second, batch-specific additive and multiplicative effects 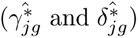 are estimated using the empirical Bayes framework, with the uncertainty of these site-specific estimates decreasing with increasing sample size. Crucially, both steps can introduce asymmetry because batches are not adjusted equally or reciprocally when their sample sizes differ. In the standardisation step, global mean and variance estimates are dominated by larger batches, causing smaller batches to be shifted and scaled more aggressively toward the pooled distribution. In the empirical Bayes step, batch-specific parameters are shrunk toward the global prior to different extents, with smaller batches undergoing stronger shrinkage than larger ones. As a result, corrections are disproportionately driven by larger batches, and the effects introduced during standardisation propagate into the subsequent parameter estimation.

Previous works that investigated ComBat’s performance have reported that successful harmonisation is usually achieved with *≈* 30 participants per batch, with different metrics being used to define success in each case (Fortin et al., 2017; Orlhac et al., 2022). Often in these cases, the statistical evaluation of the batch effect in the harmonised data is used to assess whether or not “successful” harmonisation has been achieved, but not a direct assessment of the predicted correction terms themselves. Here, as we have not only a very large sample size but also a ground-truth batch, we are able to investigate the batch effect directly across absolute batch sizes, batch proportion relative to total sample size and the overall severity of batch differences in terms of the additive and multiplicative effects. This design allows us to characterise when ComBat’s correction terms become stable and reliable, providing practical guidance on the dependence of harmonisation certainty and magnitude on sample size, batch imbalance, and batch-effect severity—factors that are especially critical in downstream machine learning applications.

To do this, we take randomly drawn participants from each batch at 100 evenly spaced sample sizes between 10 and 983, corresponding to the full size of the missing T2-FLAIR batch after outlier removal, and apply ComBat (method 5) on the selected sample, extracting the values of the ComBat corrections 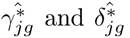. This is repeated 20 times for each sample size in order to provide a reliable estimate for the mean and variance for the ComBat correction terms across different sample sizes. To assess if the proportion of the batch has any impact on the reliability of the estimated ComBat correction, we repeated this at different ratios between the two FST2 batches, showing the mean and variance of the estimates at the following ratios between batches: 1:1, 1:2, 1:5 and finally 1:10. By varying the relative batch proportion and total sample sizes in this way, we can show empirically at which sample sizes the batch effect is well defined enough to have consistent values for its correction. We then use the previously estimated Cohen’s d and *V_r_* as indicators of the severity of the batch effect, comparing IDPs that have different severities to see how the magnitude of the batch effect changes how variable the estimated correction is, something which has not been commented on before.

## 3 Results

### 3.1 Characterising the batch effect

We first looked at the mean shift and scaling components of the batch effect by calculating Cohen’s d and the ratio of variance between the two batches for both the whole sample and the 100 within-subject repeat group (Figure 1).

**Figure 1:**
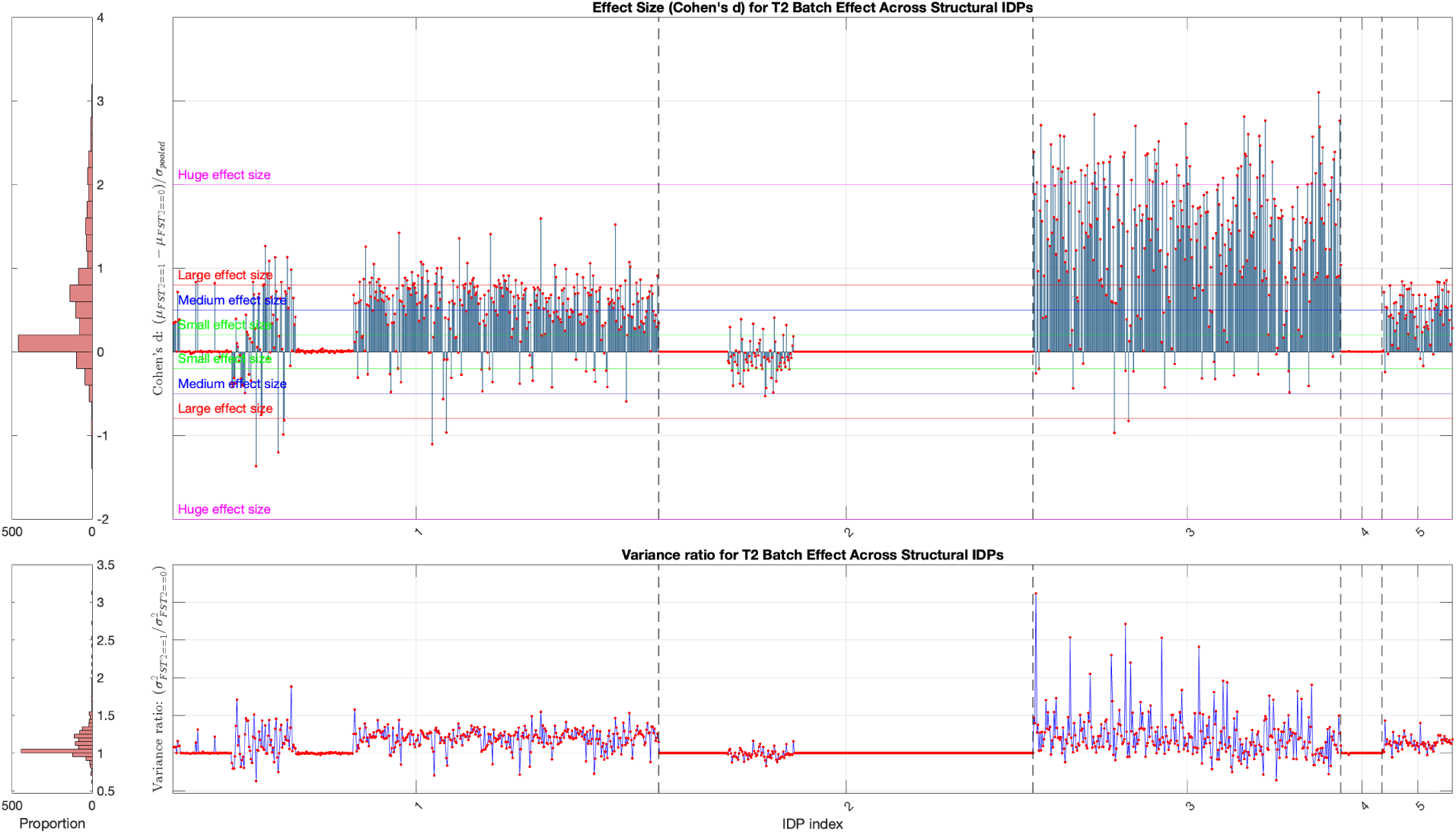
Cohen’s d (top) illustrating the batch effect on the mean and *V_r_* (bottom) showing the scaling effect between the two batches within the whole UKB sample. IDPs are grouped into the following five categories: (1) Regional and tissue volume (both cortical and subcortical), (2) cortical area, (3) cortical thickness, (4) regional and tissue intensity, and (5) grey-white matter contrast. The lines and dots in this plot, and later figures of the same format, show the same value, with the dots being included to allow for easier interpretation.

Most IDP differences (batch bias) in both the mean and variance were similar in the within-participant repeat group and the whole UKB sample; however, a small negative Cohen’s d score was observed in some IDPs in the whole sample that wasn’t present in the within-subject repeat group. This difference was due to slightly unbalanced sex and age samples between the batch with T2-FLAIR scans and without (64.6 versus 62.6 years and 55.4% female compared to 54.8% female, respectively). This was verified through regressing out the FST2 effect while preserving the other confounds and comparing the results with the within-participant group.

The largest difference in both the mean and variance was in cortical thickness IDPs (category 3), with the batch without T2-FLAIR having systematically lower estimates for the thickness across the cortex, as well as a substantially lower variance within this group. There were some additional differences in cortical volume estimates, but very little difference in cortical area. Because cortical volume depends on both cortical area and thickness, the observed volume differences are likely driven primarily by the thickness effect and that the cortical thickness measures are the most affected by the missing T2-FLAIR.

To further characterise this effect, we computed the within-subject differences (in mm) between paired measurements for 100 repeated participants. This was done for each of the 306 cortical thickness IDPs per participant, yielding 100×306 subject–IDP differences. The histograms in Figure 2 show the distribution of these differences across all subject–IDP pairs (i.e., without averaging across subjects or IDPs) for both the absolute difference in mm and the percentage difference relative to the measure obtained with T2-FLAIR.

**Figure 2:**
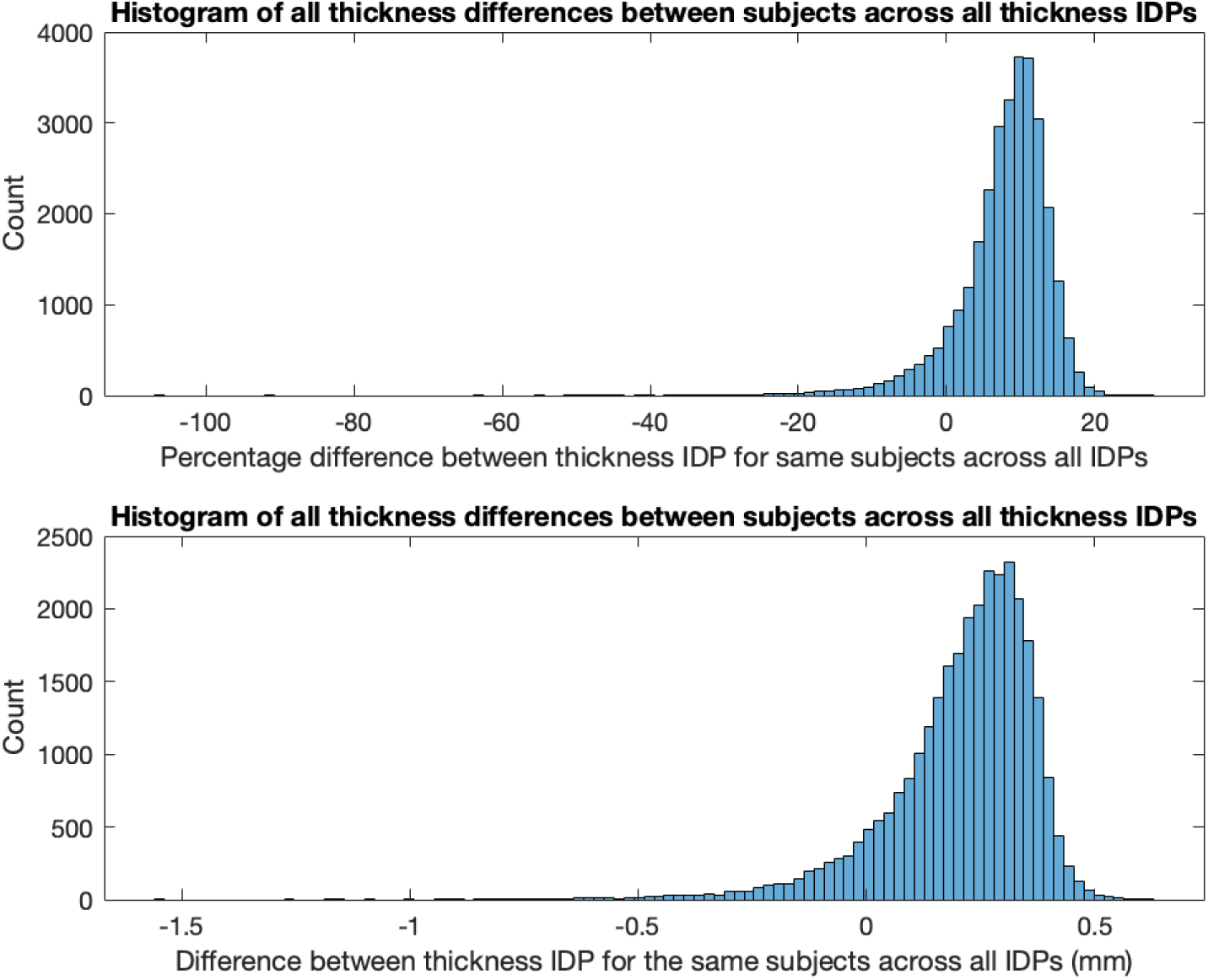
Histograms showing the percentage difference (top) and the absolute difference in mm (bottom) in the within-participant repeat measures of 306 unique cortical thickness IDPs. Here, the participant measures without T2-FLAIR used are subtracted from the measures with T2-FLAIR used, with a positive measure thus indicating a reduction when T2-FLAIR is not used.

As shown in Figure 2, it was found that the mean difference across all cortical thickness ROIs was a 0.3 mm reduction in thickness, indicating that when T2-FLAIR was not used, the thickness measures were relatively reduced. This appeared to arise from a systematic outward displacement of the pial surface estimate when T2-FLAIR was included. Additionally, including T2-FLAIR helped to correct gross pial estimation errors where the contrast of the T1 is ambiguous, leading to more accurate surface modelling. The locations where the surface fit tends to fail are those where the expected gradient between the grey matter and CSF is missing due to the signal intensity from the meninges being similar to grey matter in these regions. This systematic difference in pial surface estimation, as well as the regions which have a higher rate of failure when T2-FLAIR isn’t used, can be seen in Figure 3, where the average surfaces for 100 participants with T2-FLAIR (blue) and without T2-FLAIR (red) are overlaid.

**Figure 3:**
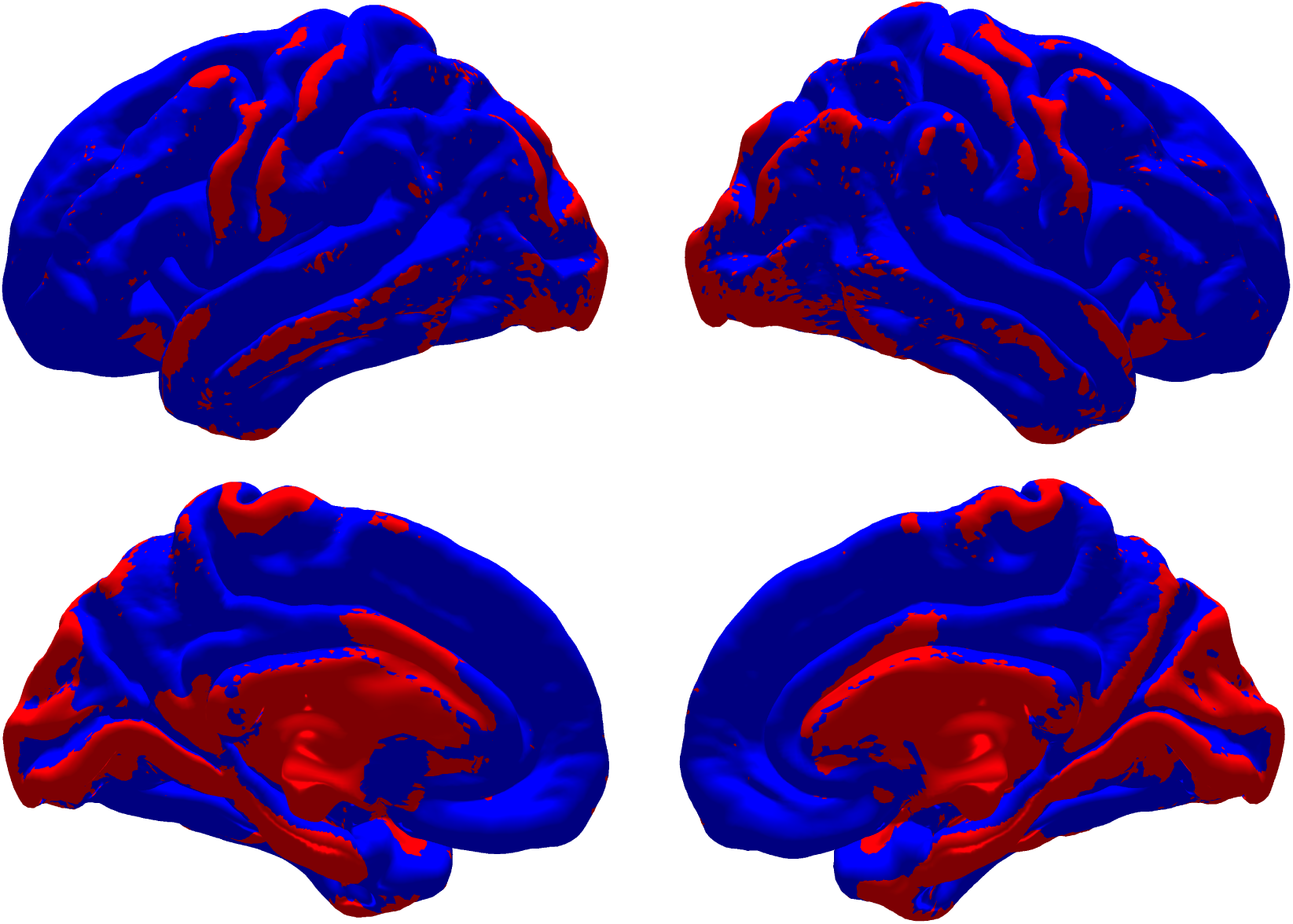
Differences in pial surface estimation with (blue) and without (red) the inclusion of T2-FLAIR volumes, averaged across 100 participants. Blue regions indicate areas where the pial surface extends further away from the white matter surface when including T2-FLAIR, and vice versa for red. The overall effect is that including T2-FLAIR leads to higher cortical thickness estimates across most of the brain (blue areas). The red areas correspond with known regions (particularly in UKB data) where the T1-weighted intensity of the meninges is very similar to that of grey matter, leading to spuriously thick cortical estimates.

Despite the bias mainly affecting IDPs of cortical thickness and volume, a small subset of cortical area IDPs also had medium effect sizes between the two batches (Figure 1). These area IDPs are those derived largely from the regions shown in red in Figure 3, for example, the parahippocampal, lingual, cuneus and entorhinal areas all showed a negative medium-to-large effect size (*< −*0.6 Cohen’s d) score, indicating relatively larger estimated surface areas in these regions when not using T2-FLAIR. An example of the surface fitting for the same individual with the T2-FLAIR image (blue) and without (red) can be seen in Figure 4. The chosen case represents a relatively severe failure case, showing an example where not using T2-FLAIR causes a failure of the gross pial surface estimation in the posterior/inferior aspect of the occipital lobe (bottom of the image).

**Figure 4:**
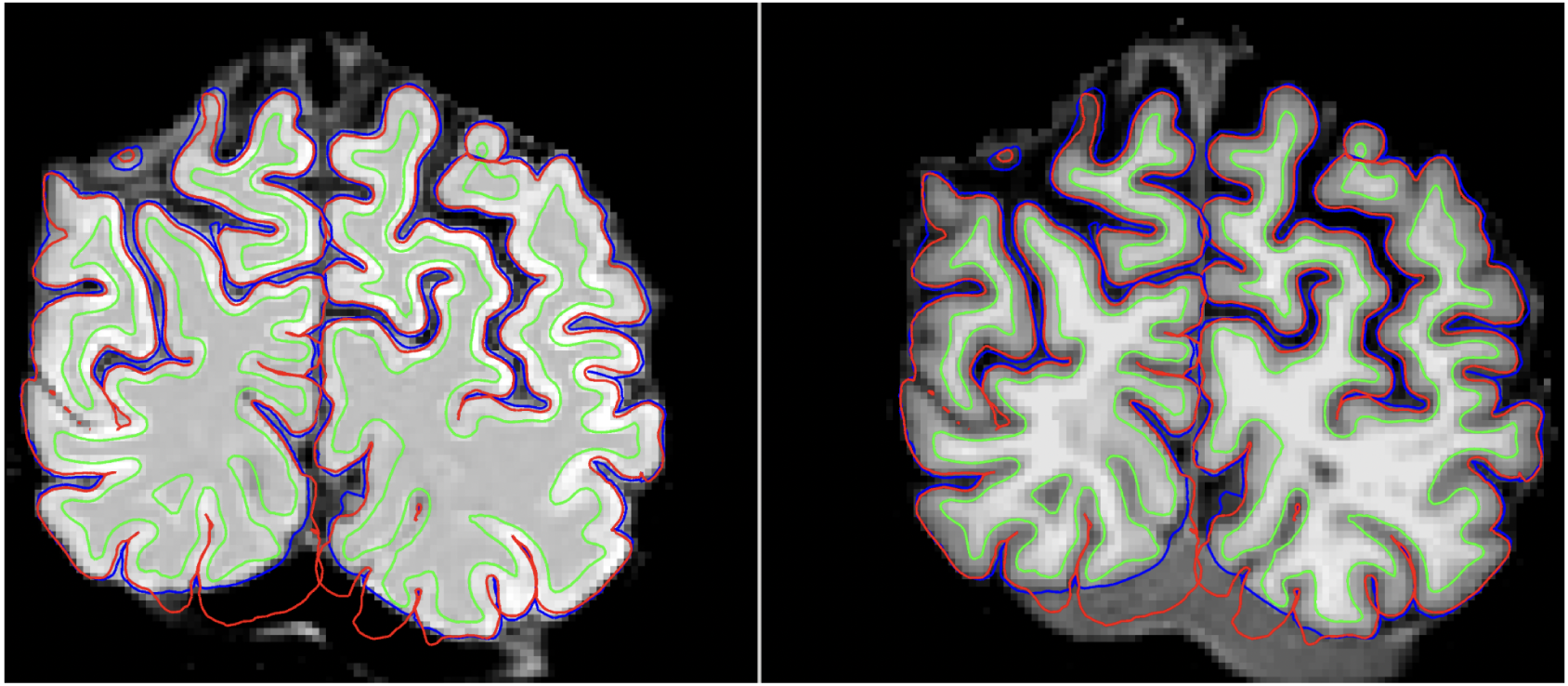
Pial surface estimates with T2-FLAIR used in FreeSurfer (blue) and without T2-FLAIR used (red) overlaid onto the T1 image (right) and the T2-FLAIR image (left) for one participant.

To assess agreement in cortical thickness estimates, we computed Pearson’s correlation coefficients between all pairs of thickness IDPs (ROIs) within the repeat participant group, separately for processing with and without T2-FLAIR. This yields a distribution of ROI–ROI correlations reflecting the covariance structure of cortical thickness estimates under each processing condition. The correlations obtained without T2-FLAIR were systematically lower than those obtained with T2-FLAIR (Figure 5), suggesting greater measurement variability and reduced internal consistency in cortical thickness estimation when T2-FLAIR is omitted. While increased inter-regional correlations do not directly imply improved biological accuracy, these findings are consistent with the observed reduction in gross pial surface estimation errors when T2-FLAIR is included.

**Figure 5:**
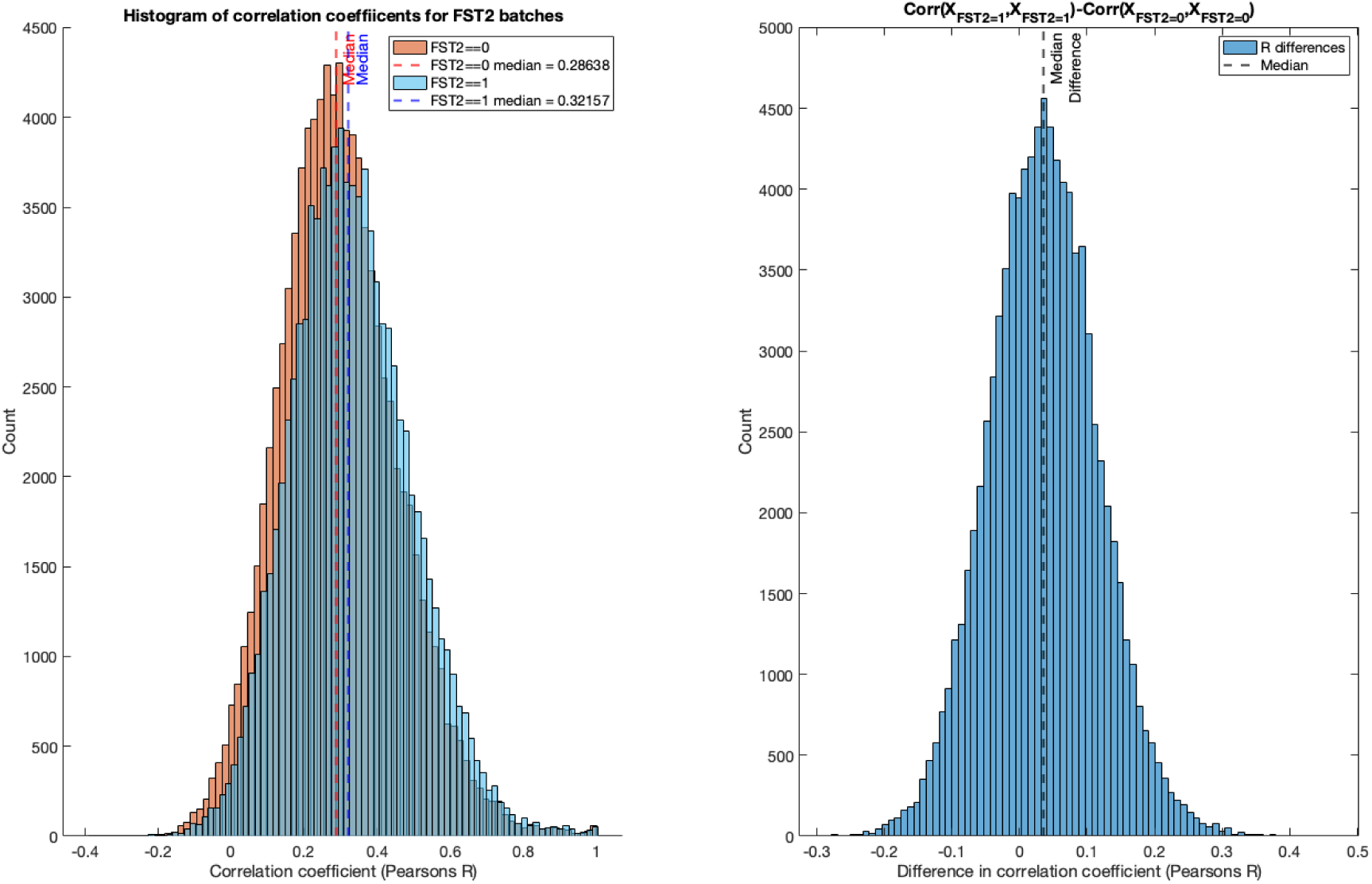
Pearson’s correlation coefficient between thickness IDPs for the same participants with (orange) and without T2-FLAIR (blue) used in FreeSurfer.

To further verify that missing T2-FLAIR was an isolated effect and not impacted by any other factor that could have caused bias in the IDPs (e.g., motion during scanning), we computed correlations between a standard set of confounds (including the FST2 batch variables) and a set of quality control IDPs (QC-IDPs) taken from the pipeline, as shown in Figure 6. This was done in the whole UKB sample, and the QC-IDPs were taken from the T1 and resting state fMRI sections of the UKB protocol, which occur before the T2-FLAIR scan is taken (S. Smith et al., 2025). We did not observe strong correlations between the FST2 batch variable and any of the QC-IDPs, suggesting that image quality or motion was not the cause of the missing T2-FLAIR in FreeSurfer or any of the resulting bias in the IDPs. Additionally, there is only a very weak correlation between the missing T2-FLAIR and all of the other explanatory variables (e.g., age or sex), suggesting that the group wasn’t systematically different from the main UKB population.

**Figure 6:**
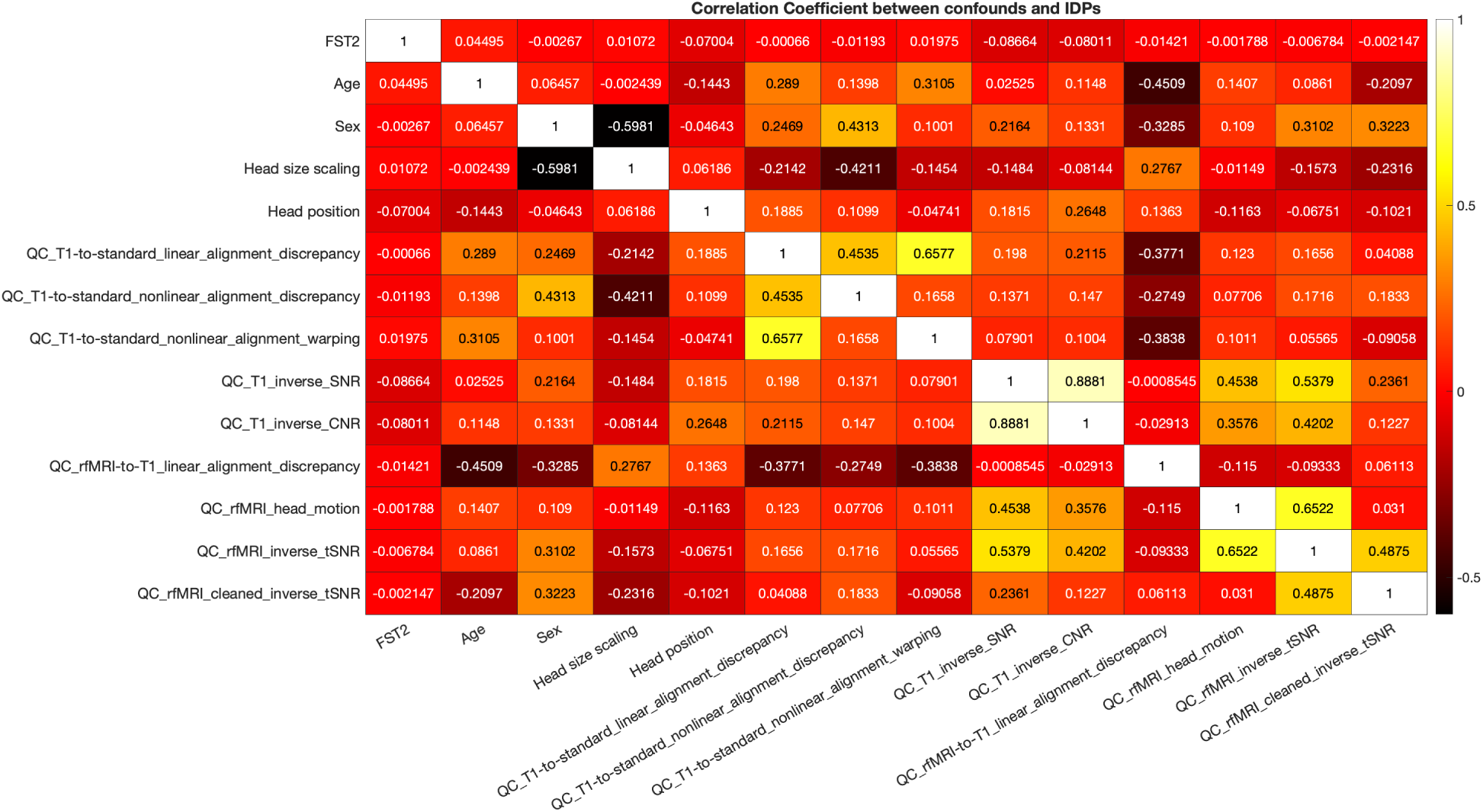
Heatmap of correlation coefficients of the set of confounds explaining the most variance against 7 quality control IDPs describing head motion and linear and non-linear alignment discrepancies between T1 and fMRI.

### 3.2 Harmonisation and deconfounding results: Removing the batch effect

After applying each of the 6 methods, we compared them against each other in terms of their ability to remove the additive component of the batch effect (Figure 7), the scaling component of the batch effect (Figure 8), and their overall ability to increase the similarity between the distributions of the two batches, using a two-sample KS-test (Figure 9). A summary of the methods can be seen in Table 1

**Figure 7:**
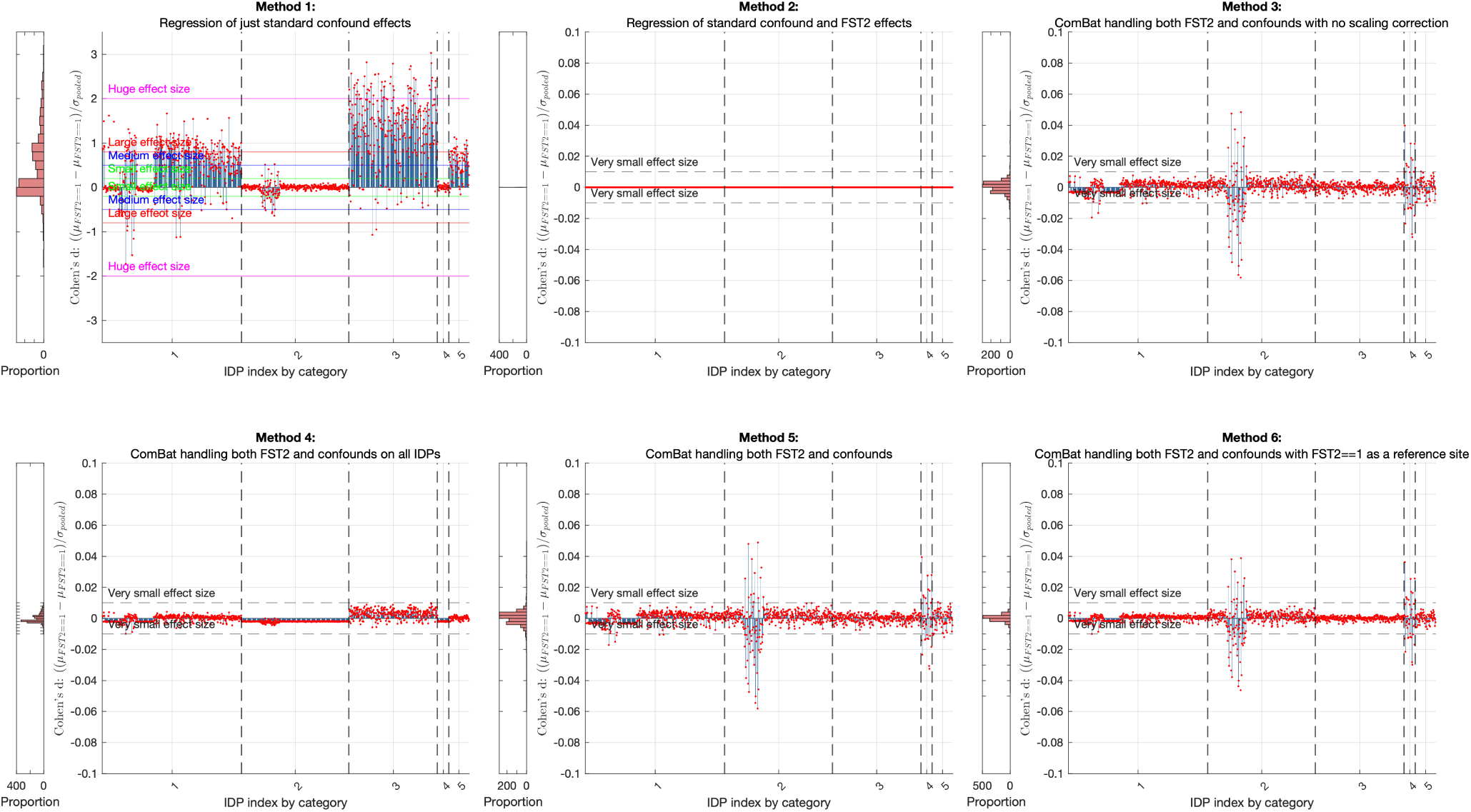
Cohen’s d for the effect on the mean for each of the methods used, plotted by IDP type, with score distribution shown on the left for each plot. The IDP categories are as follows: 1. Regional and tissue volume 2. Cortical area 3. Cortical thickness 4. Regional and tissue intensity 5. Grey-white matter contrast. Here, we include reference lines for typical magnitudes of effect sizes used in the literature.

**Figure 8:**
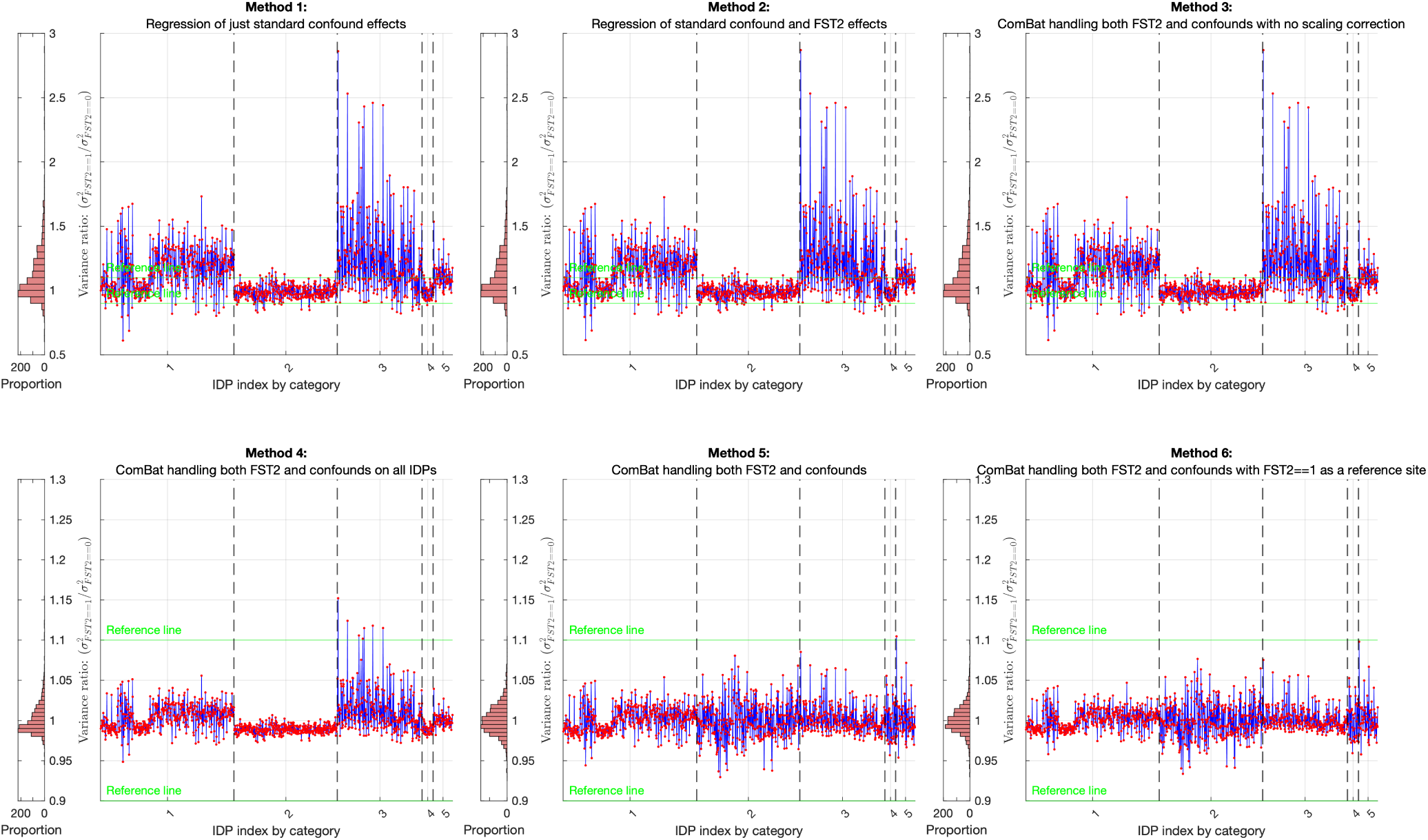
The variance ratio (*V_r_*) of the batch with T2-FLAIR vs without T2-FLAIR after applying each of the 6 methods is plotted IDP-wise for each type, with the histogram showing the distribution of variances beside the vertical axis (note the difference in axis scale).

**Figure 9:**
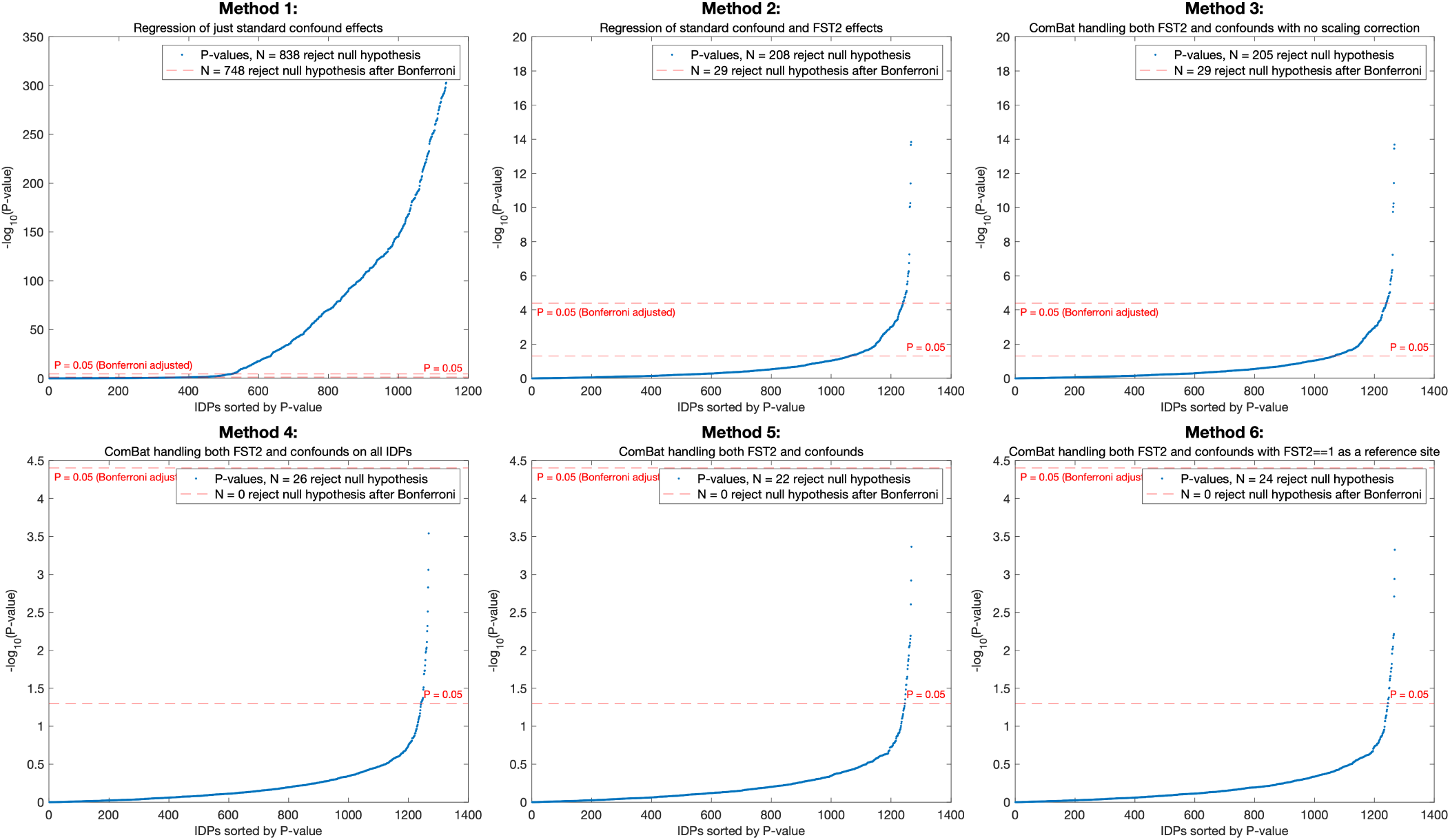
Sorted p-values from the results of a two-sample Kolmogorov-Smirnov test between the group with and without T2-FLAIR in FreeSurfer processing. Tests were performed independently for each IDP type with the null hypothesis that the two groups are from the same population.

**Table 1:** Summary of the 6 methods, showing which of them are using IDP pooling, correction on the scale, class-wise pooling and a reference batch.

| Method | Regression | Scaling correction | Prior pooling | Reference batch |
| --- | --- | --- | --- | --- |
| 1 | Standard confounds | No | No | No |
| 2 | +FST2 | No | No | No |
| 3 | ComBat additive only | No | Class-wise | No |
| 4 | Full ComBat | Yes | All classes | No |
| 5 | Full ComBat | Yes | Class-wise | No |
| 6 | Full ComBat | Yes | Class-wise | Yes |

When comparing the different methods of deconfounding and harmonisation in terms of their ability to handle the additive component of the batch effect (Figure 7), we found that all methods that directly addressed the FST2 batch effect were able to remove it. Method 1, which did not address the FST2 effect, showed a very small positive increase in the magnitude of the effect sizes when compared to no harmonisation or deconfounding. This small increase in the effect size is due to slightly unbalanced ages and sexes between the batches with and without T2-FLAIR, with this imbalance having an effect in the opposite direction to the FST2 effect (additive rather than subtractive). While the FST2 effect causes an underestimation in the thickness and volume, the slightly higher proportion of male participants and the slightly younger average age of the group missing T2-FLAIR mean that normally these measures would be higher on average. We verified this separately by modifying method 2 to instead preserve the effects of the standard set of confounds, resulting in a consistent negative shift across IDPs consistent with what we would expect.

The other five methods were able to reduce the additive batch component to well below a “small” effect size in all IDPs, with method 2 performing the best (Cohen’s d < 10^−12^ in all IDPs), showing a difference in batch means that was approximately zero. While none of the methods that used ComBat were able to reduce Cohen’s d to zero, they did all show a reduction of the effect size to well below the small effect size level (0.2). Here, method 4, which used the global pooling across IDP classes, performed better by magnitude of effect size compared to methods 3, 5 and 6 on the surface area, intensity and grey-white matter contrast IDPs (categories 2, 4 and 5). However, method 4 induced a small negative effect size systematically in the area and intensity IDPs where previously there hadn’t been one. Among the ComBat approaches, Method 4 generally performed best for IDP classes with more heterogeneous batch effects, whereas Methods 5 and 6 performed better for cortical thickness IDPs, where the batch effect was strongest and more consistent.

Figure 8 shows *V_r_* after applying each of the six methods, with reference lines at 1.1 and 0.9 in green. Methods 2 and 3 showed little improvement in *V_r_* when compared to no handling of the missing T2-FLAIR (method 1), having the same approximate mean across IDPs (1.122) and the same standard deviation of *V_r_* (0.205). Methods 4-6, which included the additional batch scaling term when modelling and correcting, showed substantial improvements. Similarly to the Cohen’s d scores between the different approaches, method 4 outperformed methods 5 and 6 in the surface area, intensity and grey-white matter contrast IDPs; performed similarly on the cortical volumes; and performed worse on the cortical thickness IDPs. The average value of *V_r_* across IDPs for methods 4, 5 and 6 were 1.0014, 1.001 and 1.001, with a respective standard deviation of 0.0188, 0.0190 and 0.0171.

The results of the two-sample KS test are shown in Figure 9. Using method 1, 838 IDPs showed significant statistical differences between the group with and without T2-FLAIR (unadjusted p-value), with 748 still significantly different at the Bonferroni-corrected significance level (*α* = 3.94*×*10^−5^). Methods 2 and 3 were respectively able to reduce this number to 208 and 207 IDPs at the unadjusted level and 29 at the Bonferroni-corrected significance levels, respectively. Methods 4, 5, and 6, which all corrected for the scaling differences between batches, performed substantially better, with each having fewer than 27 IDPs with significant differences between groups at the unadjusted p-value and zero IDPs with significant differences after correcting for multiple comparisons.

### 3.3 Mechanistic interpretation of ComBat behaviour: Prior distributions and feature-wise likelihoods

While the previous section demonstrates that both pooled (Method 4) and class-wise (Methods 3, 5, and 6) implementations of ComBat effectively reduce batch effects at the level of the chosen summary statistics, these results do not directly explain how differences in the underlying empirical Bayes formulation influence the estimated batch corrections.

In particular, although the class-wise implementation is expected to produce more informative priors—due to being estimated from more homogeneous feature sets—this did not necessarily translate into improved performance in summary statistics such as Cohen’s d or variance ratio.

To better understand the behaviour of these methods, we examine the prior distributions governing the batch-specific parameters and the corresponding feature-wise likelihoods (Figure 10). In particular, we compare the globally pooled implementation (Method 4), in which all IDPs contribute to a shared prior, with the standard class-wise implementation (Method 5), where priors are estimated separately within each IDP category. This allows us to directly assess how prior structure influences shrinkage and the resulting parameter estimates.

**Figure 10:**
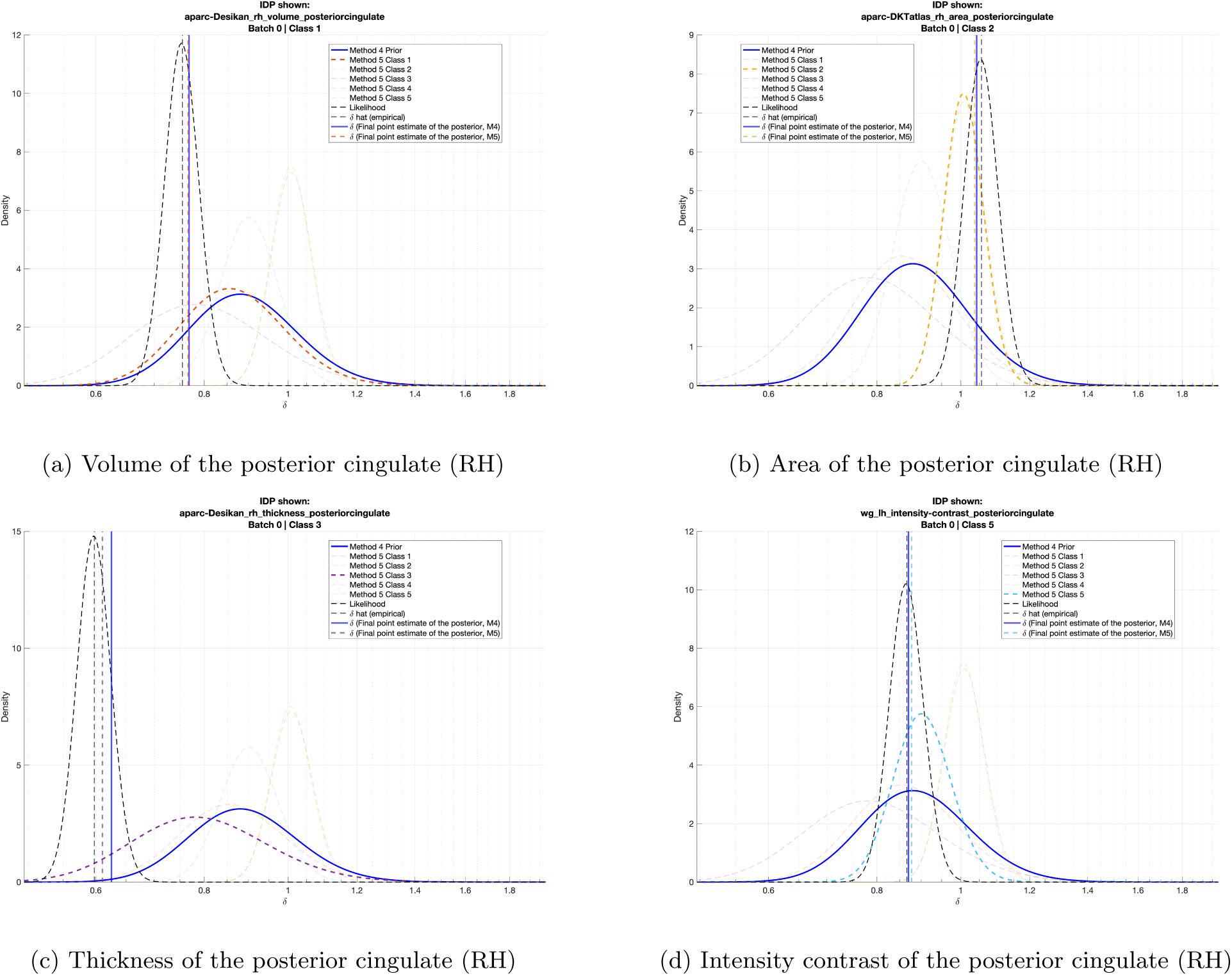
Empirical prior distributions for each of the categories in method 5 and the single global prior used in method 4 for *δ*^2^, plotted with the respective final point estimate for the correction on the variance and the likelihood of the initial estimate of the correction before applying the iterative empirical Bayes shrinkage. For each IDP, the class-wise prior distributions are faded if they do not belong to the same class as the final IDP-specific point estimate being plotted. We show here 4 IDPs from the batch that did not have the T2-FLAIR used in FreeSurfer, showing, respectively, the predicted correction on the scale of the Volume (a), Area (b), Thickness (c) and Intensity Contrast (d) of the posterior cingulate.

Here, we focus on Methods 4 and 5 because they share the same initial feature-wise estimates of the batch effect (i.e., maximum likelihood estimates) and therefore the same likelihood. As such, any differences in the final parameter estimates arise solely from differences in the empirical Bayes priors. Comparing the empirical priors between methods 4 and 5 against the final point estimate of the correction of *δ*^∗^ allows us to see how the global pooling of method 4 compared to the class-specific pooling of the other ComBat approaches. In each case, the initial likelihood function for the first estimate of *δ*^∗^ is displayed. We showed it in this way, as each subsequent likelihood is estimated from the residuals after subtracting the predicted mean effect *γ*^∗^, meaning methods 4 and 5 will have different subsequent likelihoods after the first ComBat iteration.

Figure 10 illustrates how the choice of prior pooling strategy influences the final variance-scaling estimates across IDP classes. For the posterior cingulate volume IDP (panel a), the global and class-wise priors were similar in both location and spread, resulting in closely aligned final estimates. For cortical area (panel b), the class-wise prior was narrower and more closely aligned with the feature-wise likelihood than the global prior, supporting stronger class-specific shrinkage. In contrast, for cortical thickness (panel c), the global prior was slightly tighter but shifted further from the likelihood, consistent with the poorer scaling performance of Method 4 for the IDP class most affected by the FST2 batch effect. For grey-white matter contrast (panel d), the global prior lay closer to the likelihood than the class-wise prior, matching the stronger performance of Method 4 for this IDP category. These examples show that the relationship between the empirical prior, feature-wise likelihood, and final point estimate differs across IDP classes, with the largest visible differences occurring in classes where Methods 4 and 5 showed different scaling behaviours.

### 3.4 Harmonisation and deconfounding results: Correlation with non-imaging variables

We correlated the IDPs after applying each of the 6 methods against the nIDPs, which have been similarly residualised using the same set of confounds. Performing Fisher’s Z-transform on the correlations allows us to express the results as BA plots of agreement between each method, assessing the direct impact of each additional term from ComBat and each of the variations in applying ComBat (methods 4-6).

When comparing methods 2 and 3 against method 1 (removal of only standard confound effects), the vast majority of correlations were in agreement between the three approaches, showing a preservation of correlations when addressing the additive component of the FST2 effect (Figure 11). Additionally, strong negative correlations between volume measures and bone mineral density were strengthened when the FST2 effect was removed, implying that the FST2 effect could dampen real, strong biological associations if not corrected for. While the vast majority of correlations showed agreement, a small minority had larger differences, as observed by the positive and negative tails centred around 0. These tails are composed of the correlations of cortical thickness IDPs and health-related nIDPs, with the largest difference being multiple IDPs with caffeine consumption on the day of the scan, ICD-10 coded values describing unspecified disturbances of taste or smell (Stroganov et al., 2022; Sudlow et al., 2015). Through individual inspection, the largest differences showed stronger correlations with method 1 compared to methods 2 and 3, indicating that these correlations could be driven by the FST2 batch effect rather than actual correlations of brain measures with these non-imaging variables.

**Figure 11:**
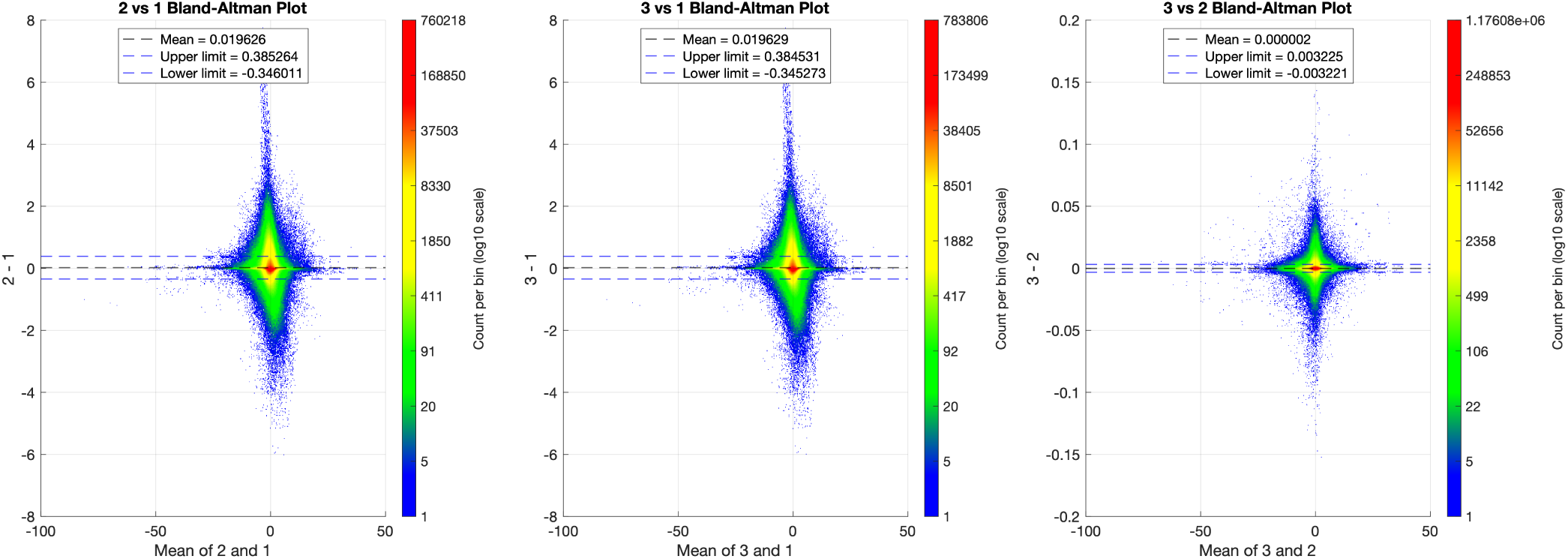
Bland-Altman plots of agreement between the first 3 methods for deconfounding and batch correcting. 2 vs 1: Residualising using just confounds against using confounds plus the binary FST2 vector. 3 vs 1: Residualising using just confounds against using ComBat for deconfounding and batch correction with no variance correction. 3 vs 2: Residualising using confounds plus the binary FST2 vector against using ComBat for deconfounding and batch correction with no variance correction.

When comparing the different ComBat approaches (methods 4-6), we can see the differences in correlation are far lower than those observed between method 1 and methods 2 and 3 (note the difference in y-axis limits). Between the ComBat approaches, the largest difference was the addition of the scaling term (Figure 12, left panel), with method 3 being in closer agreement with method 2 than method 5 (see Figure 11). Applying ComBat on distinct categories of IDPs (method 5) had very minor differences when compared to applying ComBat to all IDPs regardless of category (method 4), with the differences being less than 0.15 for all IDP-nIDP pairs. ComBat applied with and without a reference batch (methods 5 and 6) showed the greatest agreement between correlations, with the differences being normally distributed between -0.04 and +0.04 with an overall mean difference of approximately 0. This high level of agreement between methods is expected, as even without using a reference batch, the number of participants in the batch with T2-FLAIR is so much greater than those without (43,470 vs 983) that it dominates estimates of the mean and variance of the data, resulting in the standardisation step within ComBat being heavily biased towards the larger batch.

**Figure 12:**
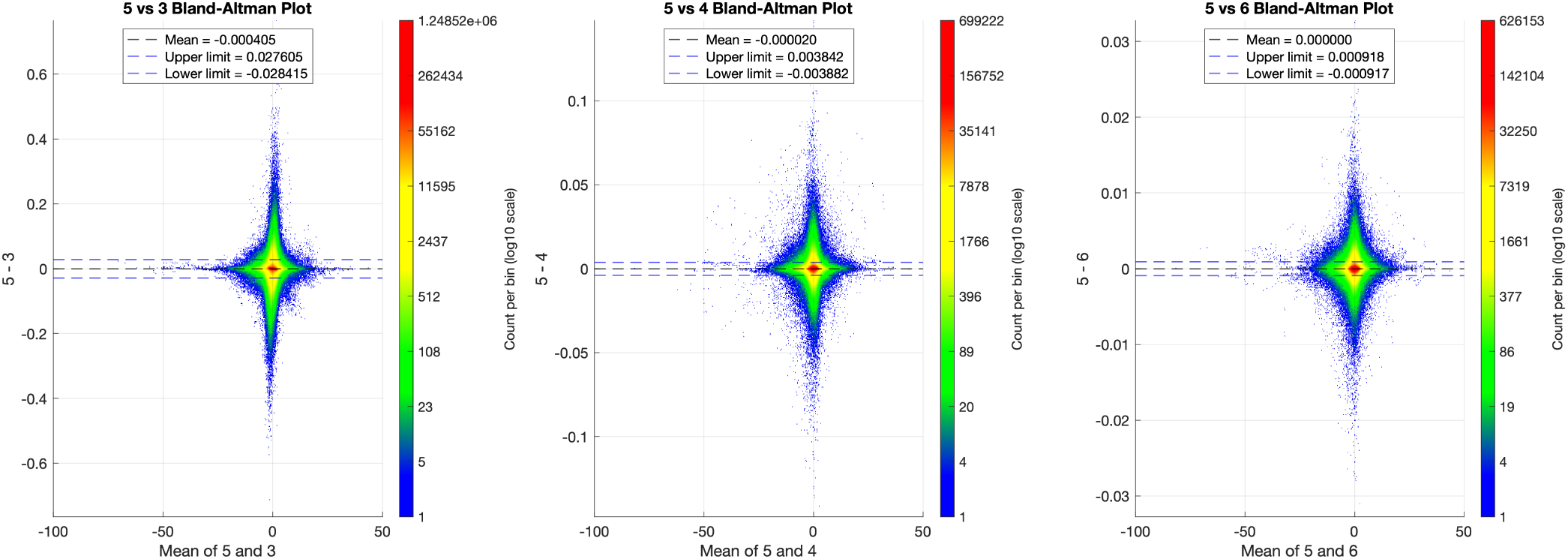
Bland-Altman plots of agreement between the different ComBat approaches (methods 3-6). 5 vs 3: ComBat applied class-wise, with the additional scaling term omitted vs ComBat applied class-wise with the scaling term included. 5 vs 4: ComBat applied class-wise vs ComBat used on all IDPs at once. 5 vs 6: ComBat applied class-wise using no reference batch vs ComBat applied class-wise using the FST2=1 batch as a reference.

**Figure 13:**
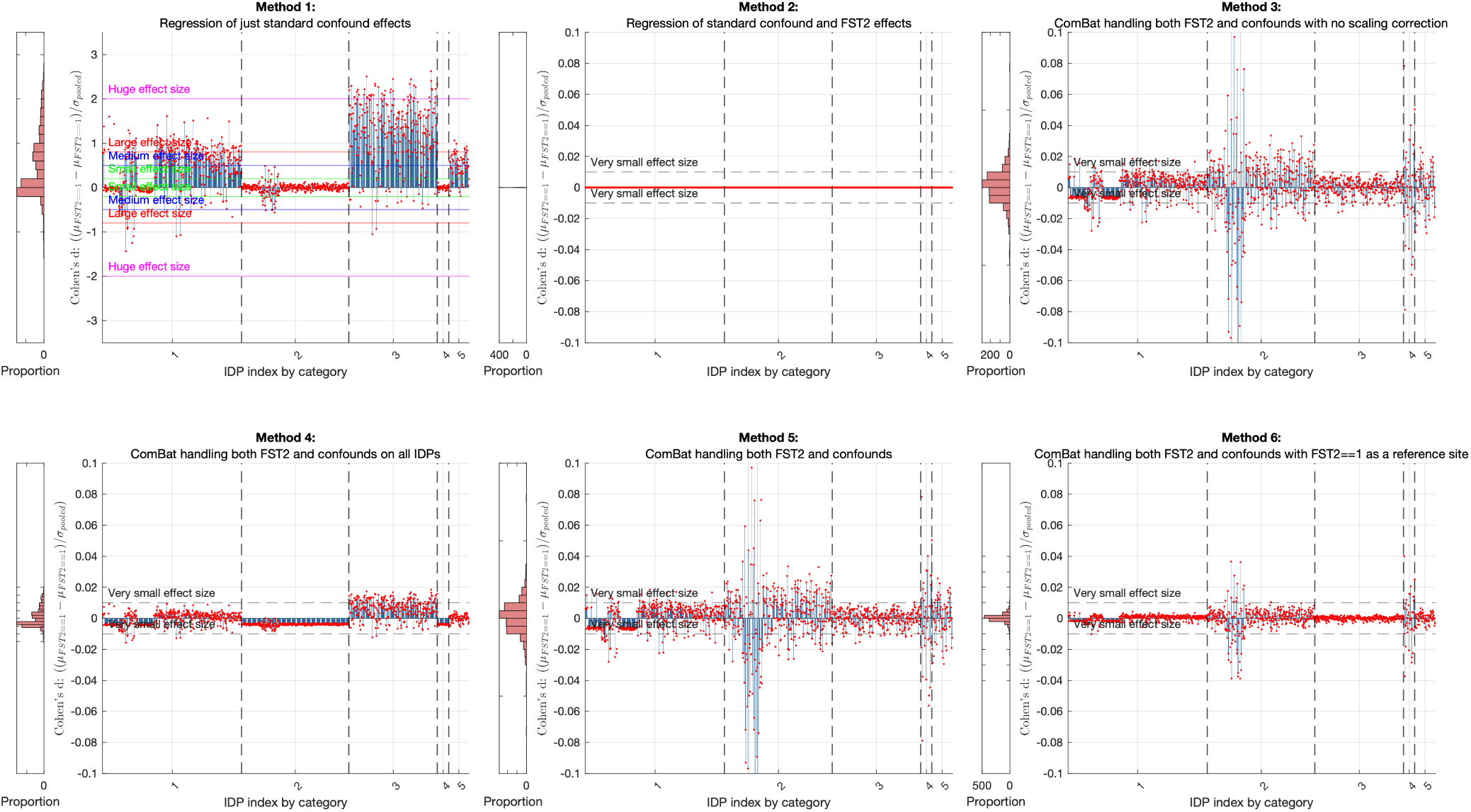
Cohen’s d for the effect of batch on the mean for each of the methods used when batch proportions are equal, plotted by IDP category, with the distribution shown on the left for each plot.

### 3.5 ComBat changes with sample size and batch proportions

#### 3.5.1 Removing the batch effect

To test how harmonisation efficacy changed at different batch proportions and sizes, we first took a random subsample of the larger batch with T2-FLAIR at the same sample size as the smaller batch (N = 983 after outlier removal) and reran each of our methods on this new dataset. We then computed Cohen’s d, *V_r_*, and the results of the two-sample KS test as we had done before.

When comparing Cohen’s d, all methods that addressed the FST2 mean component were still able to reduce the mean shift bias to below a small effect size (0.2) (Sawilowsky, 2009). Methods 3 and 5 showed significantly larger Cohen’s d scores, particularly in the area IDPs discussed in section 3.1, for which the largest difference was seen. Method 4 performed slightly worse when the unbalanced sample sizes were used. In all but area and intensity IDPs, method 6 showed the best performance, with the best reduction of effect size in the cortical volume and thickness IDPs.

A similar result was seen with the ratio of variance (Figure 14), with methods 1, 2, and 3 still offering little improvement, in fact increasing the difference in some IDPs. Method 4 improved the ratio of variance between batches substantially, but not as well as when the unbalanced full dataset was used. While method 4 was able to reduce *V_r_*, it performed worse than when the unbalanced sample was used, particularly in the thickness IDPs where the variability is most severe. Method 5 did slightly better than method 4 in the thickness IDPs, but comparatively worse in other classes. Method 6 performed the best here in all IDPs, with no IDP having a *V_r_* greater than 1.1 or less than 0.9. The mean ratio of variance of the 3 latter methods were 1.001, 1.001 and 1.002, with standard deviations being higher for methods 4 and 5 (0.04 and 0.041) compared to method 6 (0.016).

**Figure 14:**
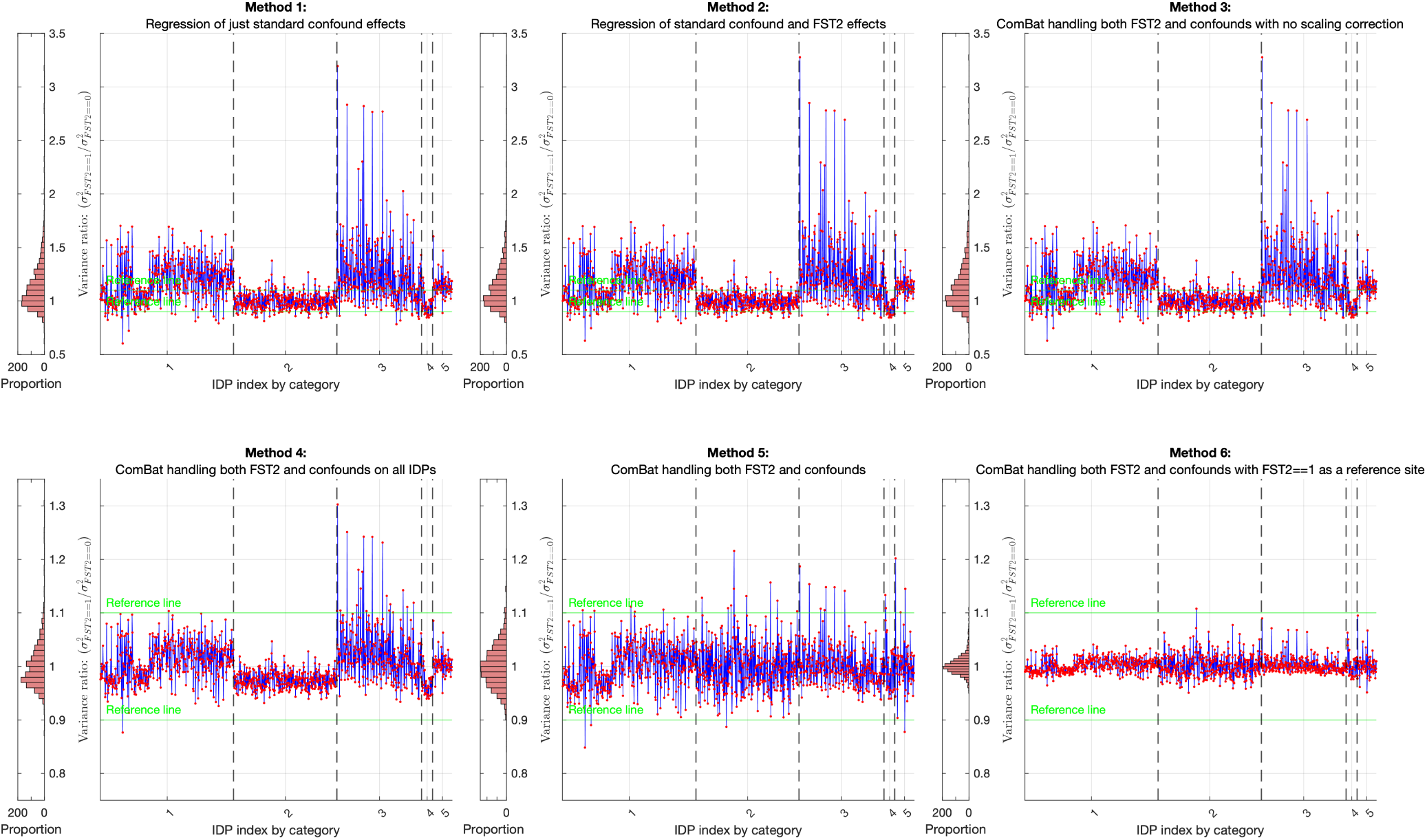
Variance ratio (*V_r_*) between batches after applying each of the six methods when batch proportions are equal, plotted IDP-wise for each category, with the histogram showing the distribution of variance ratios in each case.

For the results of the two-sample KS tests, methods 2 and 3 rejected the null hypothesis less frequently than when the full dataset was used, with methods 4, 5, and 6 also performing better on average (Figure 15). This is because the critical value of *D_m,n_* required for statistical significance decreases with increasing sample size, making the KS test more sensitive for larger samples. Consequently, smaller samples reduce the test’s ability to detect differences between distributions.

**Figure 15:**
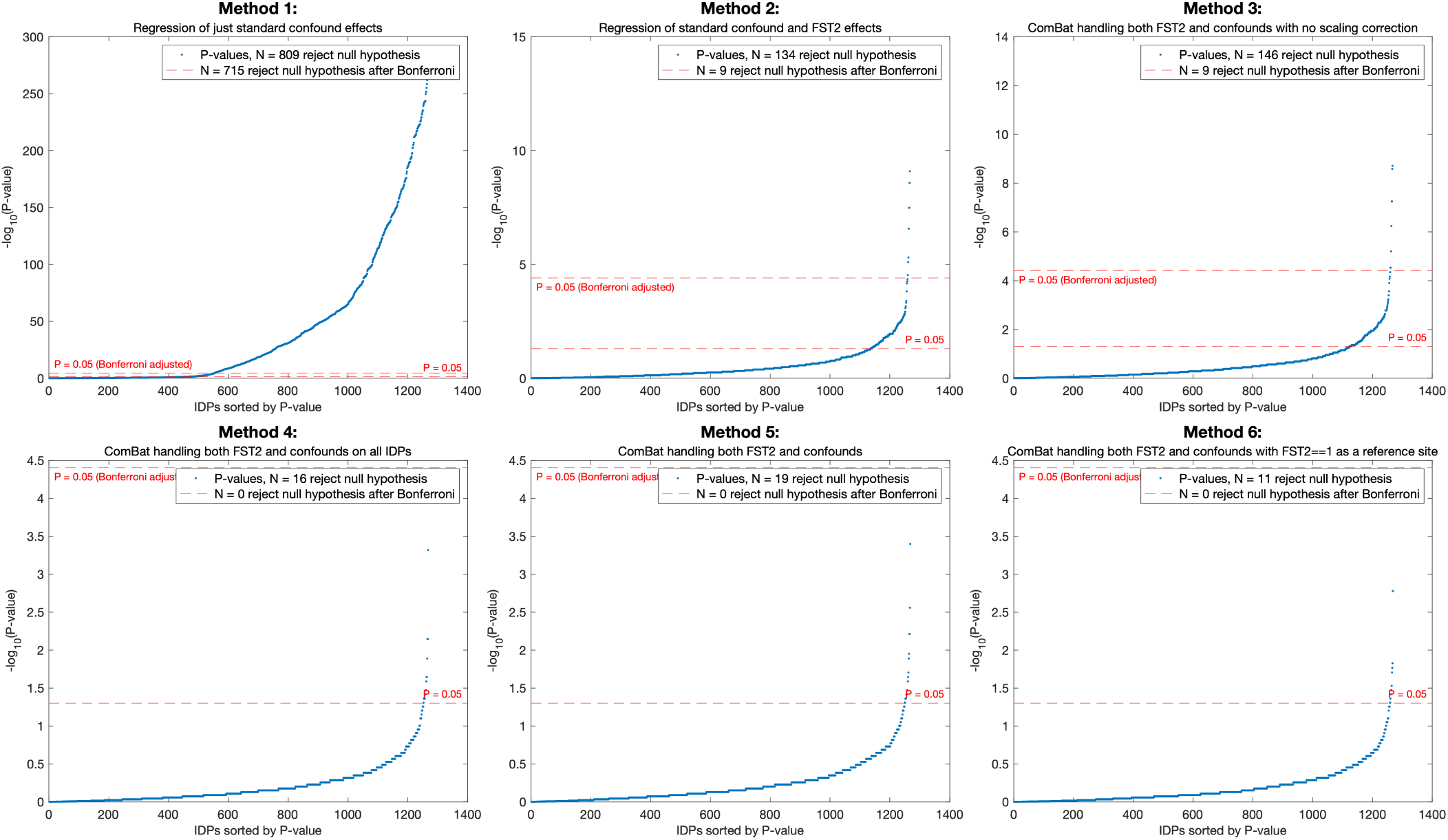
Sorted p-values from the results of a two-sample Kolmogorov-Smirnov test between the group with and without T2-FLAIR in FreeSurfer processing. Tests were performed independently for each IDP type with the null hypothesis that the two groups are from the same population.

#### 3.5.2 Correlation with non-imaging variables

We next examined whether the preservation of IDP–nIDP correlations differed when the two FST2 batches were represented equally. As in the full-sample analysis, correlations were Fisher-Z transformed and compared using Bland-Altman plots across all IDP–nIDP pairs. The comparisons between methods 1, 2 and 3 in the equal batch-size dataset showed a pattern highly similar to that observed in the full, unbalanced sample (Figure 16). Where Methods 2 and 3 differ from 1, they generally show a weaker association between IDPs and nIDPs, regardless of the sign of the correlation. This most likely shows that the methods are removing shared confounds. Methods 2 and 3 are very similar to each other. This indicates that, even when batch sizes were balanced, explicitly modelling the FST2 effect altered a subset of correlations in a consistent manner, whereas the empirical-Bayes treatment of the additive effect in method 3 produced results very similar to standard regression-based correction.

**Figure 16:**
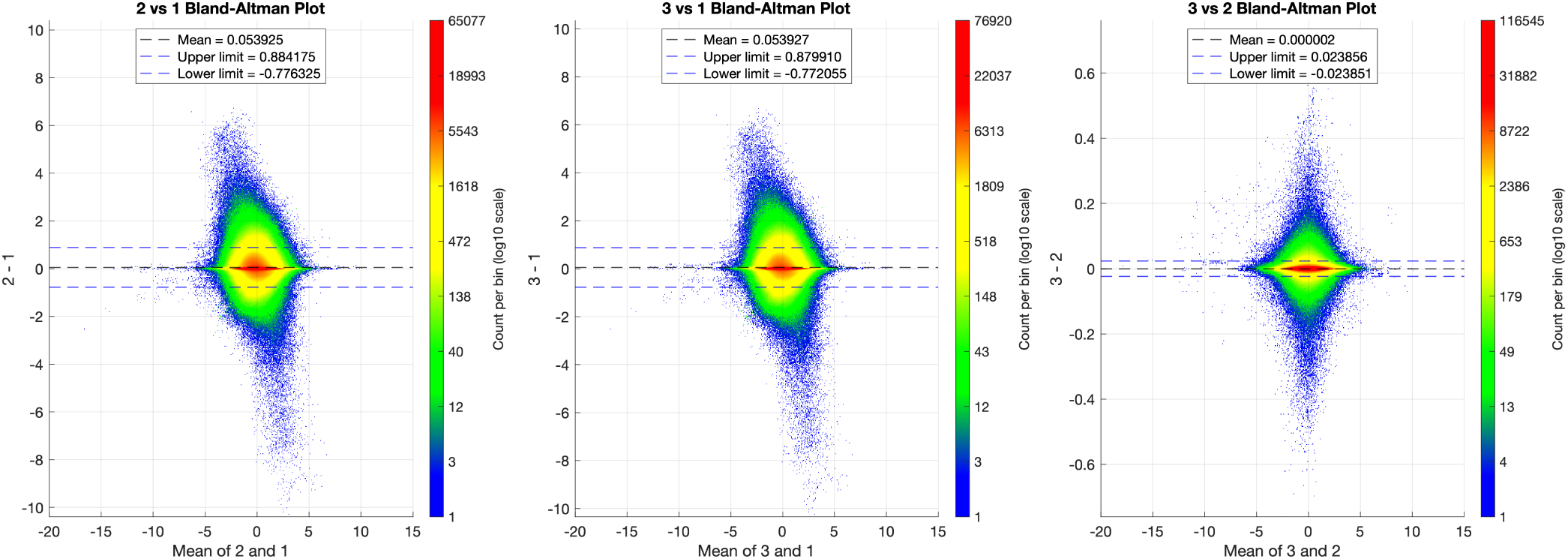
Bland-Altman plots of agreement between the first three methods in the equal batch-size dataset. 2 vs 1: residualising using standard confounds only compared with residualising using standard confounds plus the binary FST2 vector. 3 vs 1: residualising using standard confounds only compared with ComBat-based correction without variance correction. 3 vs 2: residualising using standard confounds plus the binary FST2 vector compared with ComBat-based correction without variance correction.

When comparing the ComBat approaches in the equal batch-size dataset, larger differences were observed than in the full unbalanced sample, particularly for the comparison between method 5 and method 3 (Figure 17). This comparison isolates the effect of including the ComBat scaling term and shows wider limits of agreement than the comparisons between the full ComBat approaches. This suggests that variance correction has a measurable effect on downstream IDP–nIDP correlations when the two batches contribute equally to the estimation of the data distribution. In contrast, methods 4 and 5 showed very close agreement, with a mean difference close to zero and narrow limits of agreement, indicating that applying ComBat class-wise or across all IDPs jointly had little impact on downstream correlations in the balanced sample. Methods 5 and 6 also showed very strong agreement, although the small negative mean difference indicates marginally higher correlations after reference-batch ComBat. Overall, the 1:1 results support the findings from the full sample: addressing the FST2 effect changes a subset of correlations relative to standard deconfounding alone, while the different full ComBat implementations preserve broadly similar IDP–nIDP association structure.

**Figure 17:**
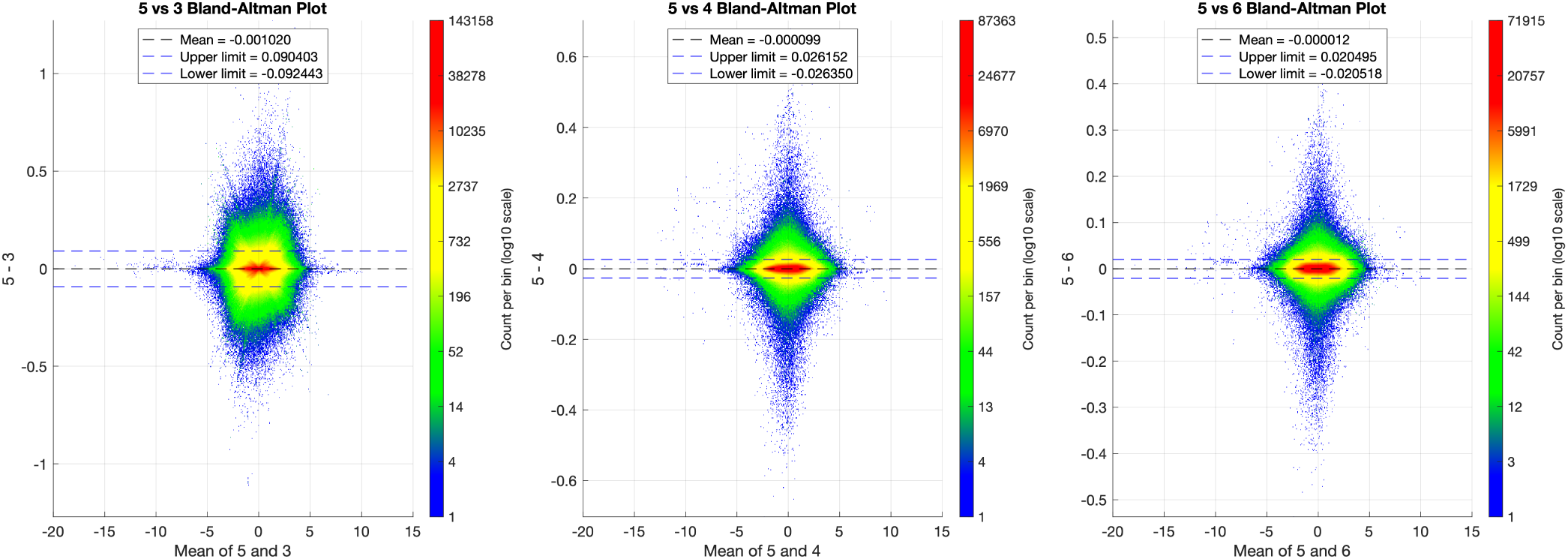
Bland-Altman plots of agreement between ComBat approaches in the equal batch-size dataset. 5 vs 3: class-wise ComBat with scaling correction compared with class-wise ComBat without scaling correction. 5 vs 4: class-wise ComBat compared with ComBat applied across all IDPs jointly. 5 vs 6: class-wise ComBat without a reference batch compared with class-wise ComBat using the FST2=1 batch as a reference.

### 3.6 Comparing ComBat adjustment terms over total and relative sample sizes and across severity of batch difference

The ComBat adjustment terms for the estimated additive batch effect 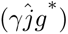 and the multiplicative batch effect 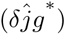 from method 5 are plotted against batch size in Figures 18 and 19, respectively. The top horizontal axis for each subplot shows the sample size of the FST2=0 batch, and the bottom horizontal axis shows the sample size for the FST2=1 batch. Each subplot shows a different total sample size but also a relative sample size, with the ratio of participants being kept constant across sample sizes for each subplot and varying from 1:1, 1:2, 1:5, and finally 1:10 (FST2=0:FST2=1) in descending order. Additionally, we show results here for two IDPs from the same area of the brain, one with a relatively large batch difference and one with a relatively minor difference, as calculated through the Cohen’s d and ratio of variance of the whole dataset. The first, displayed on the left of Figures 18 and 19, is the thickness of the posterior cingulate in the right hemisphere (Cohen’s d = 1.22, *V_r_* = 1.72). The second is the area of the posterior cingulate in the right hemisphere (Cohen’s d = -0.058, *V_r_* = 1.034). The results are displayed as the raw estimates for the additive (*γ_jg_*) and multiplicative *δ_jg_* batch terms, with the error bars being the standard deviation of the 20 repeated measures at each sample size.

**Figure 18:**
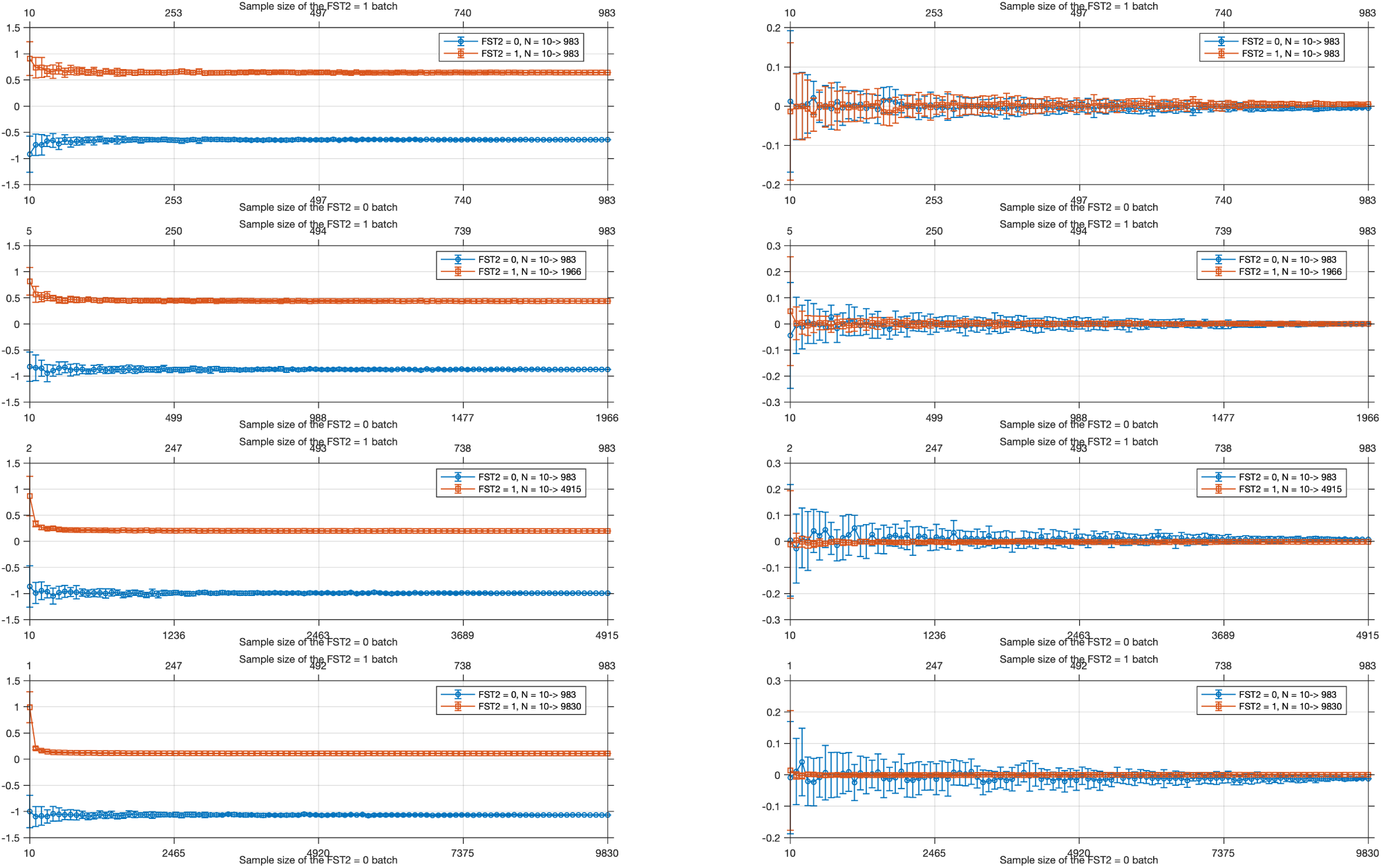
The mean value of the ComBat mean shift term *γ_jg_* is plotted over different sample sizes of the FST2=0 and FST2=1 batches at different total sample sizes and ratios between batches, with error bars showing the standard deviation of each estimate for each sample size, taken from 20 repeat measures, respectively. The left-hand plot is for the thickness of the posterior cingulate in the right hemisphere; the plot on the right is for the area of the posterior cingulate in the right hemisphere.

**Figure 19:**
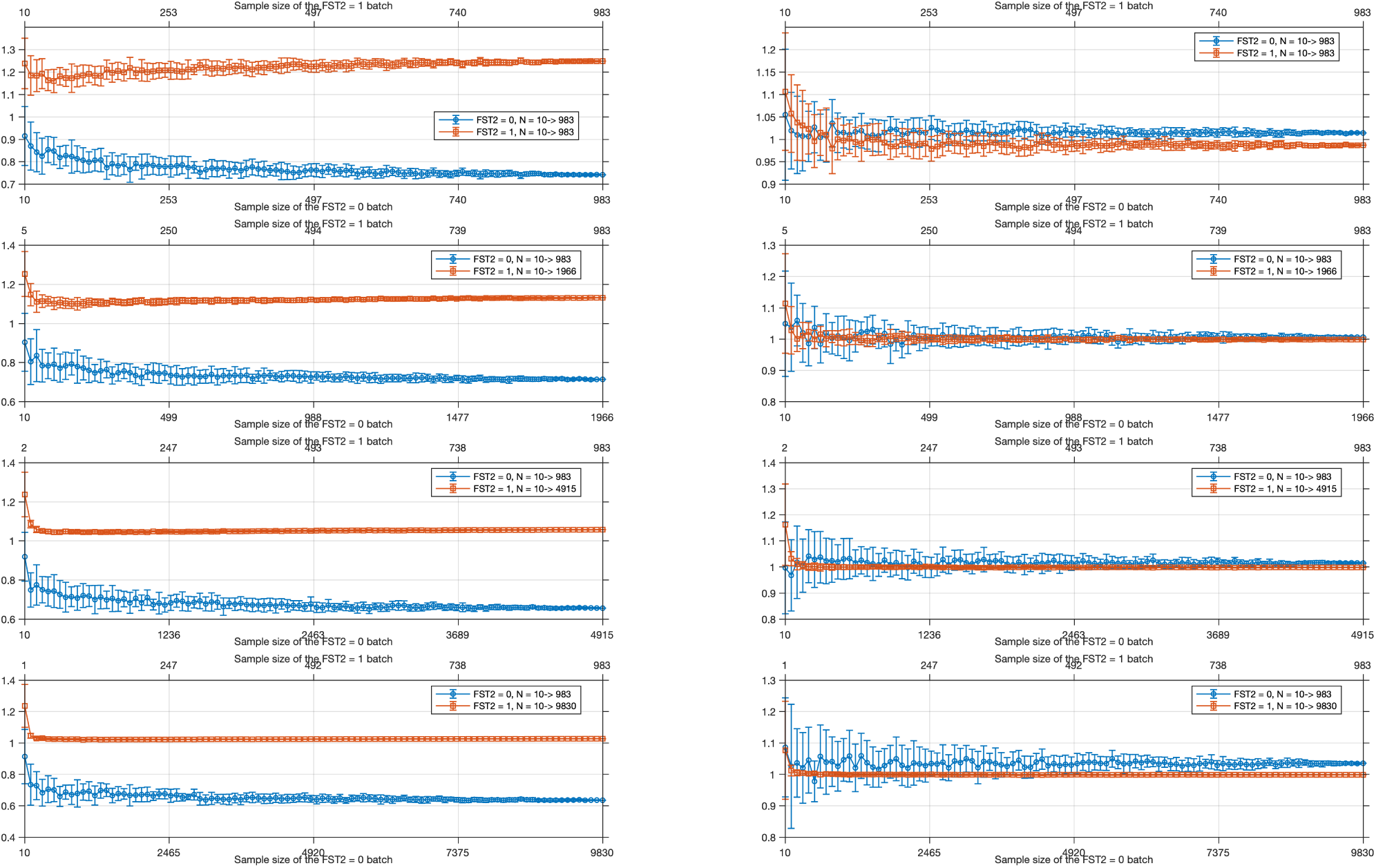
The mean value of the ComBat multiplicative term *δ_jg_* plotted over different sample sizes of the FST2=0 and FST2=1 batches at different total sample sizes and ratios between batches, with error bars showing the standard deviation of each estimate for each sample size, taken from 20 repeat measures, respectively. The left-hand plot is for the thickness of the posterior cingulate in the right hemisphere; the plot on the right is for the area of the posterior cingulate in the right hemisphere.

As shown in Figure 18, the additive component of the batch effect asymptoted fairly rapidly when plotted against sample size, with the mean estimate for the mean shift at each sample becoming consistent at around 30 samples, as is consistent with the literature (Fortin et al., 2017; Jodoin et al., 2025; Orlhac et al., 2022). Interestingly, when the batch effect was relatively small, the variability of the estimates for the mean shift at a given sample size was comparable in magnitude to the estimates for a heavily affected IDP (as shown by the magnitude of the error bars for a heavily affected IDP on the left and a less affected IDP on the right, noting the different scales of the axis).

When looking at the estimates for the scaling term in Figure 19, substantially more variation with sample size is observed than in the additive term. The leftmost plots show that when the scaling difference is large, even when averaged for multiple repeats at increasing sample sizes, the estimate for the scaling correction does not clearly approach an asymptote. Additionally, it is extremely variable with sample size, especially within the 20 to 200 sample range. When looking at an IDP with only a very minor scaling difference, we find that the values do tend to asymptote similarly to the additive term, still showing quite high variability between 10 and 200 samples, but arriving at a similar estimate for the correction at higher sample sizes. Overall, if the scaling difference between the batches is severe, the estimates for the adjustment terms are highly variable with batch size, requiring far higher sample sizes to reach a stable estimate than when the scaling difference is less severe or when compared to the additive effect.

A key observation is the asymmetry between the additive and multiplicative correction terms: while the additive term stabilises rapidly with increasing sample size (*≈* 30 subjects), the multiplicative term remains highly variable even at moderate sample sizes (*≈* 100–300), particularly for IDPs with large variance differences. This indicates that achieving stable harmonisation of scaling differences requires substantially larger datasets than those typically considered sufficient for successful ComBat application. We further verified this using the within-participant data, where sufficient mean reduction could be achieved using ComBat but *V_r_* was not substantially improved.

## 4 Discussion

This study had several aims; the first was to systematically evaluate the impact of the missing T2-FLAIR scan on the derived FreeSurfer IDPs. The second was to use different approaches for harmonising the data by treating the missing T2-FLAIR as a batch effect, producing a set of batch-corrected IDPs and showing in a sequential manner the differences between standard linear modelling and ComBat. Finally, we sought to use the large sample size and “ground truth” batch to provide an assessment of the variability of the estimated ComBat correction terms against total sample size, relative batch size and the severity of batch differences.

### 4.1 Effect of not including T2-FLAIR in FreeSurfer

The reasons for missing T2-FLAIR in FreeSurfer were varied and confirmed not to be heavily biased by age, sex, motion, site, or any of the flagged quality control parameters. Additionally, we found no strong correlations of the FST2 bias effect with any other potential confounding variables available to us (Alfaro-Almagro et al., 2021).

Excluding T2-FLAIR from FreeSurfer processing results in a systematic underestimation of the distance between the white matter surface and the pial surface estimates, with its inclusion resulting in more accurate surface fitting in regions where there were ambiguous contrast differences between grey matter and meninges in the T1 images. In some extreme cases, its exclusion can cause the pial surface fitting to fail entirely, particularly in the posterior/inferior aspect of the occipital lobe, with these failures often being missed by automatic quality control checks (Klapwijk et al., 2019). While pial fitting failures have been identified before, previous works discussing it have not considered potential downstream effects on the IDPs or evaluated the systematic underestimation of the cortical thickness that we saw here (Fischl, 2012; Lindroth et al., 2019). Contrary to our hypothesis that these inaccuracies may lead to more variable IDP measures, the examination of the within-batch variance revealed instead that the results were more affected by a systematic underestimation of the surface rather than occasional failures in surface estimation. The inclusion of T2-FLAIR in FreeSurfer processing visually improves the pial surface fitting, and omission will result in a systematic (mean 0.3 mm) underestimation of the resulting thickness IDPs.

Our results show that including T2-FLAIR improves pial surface estimation at the individual level, with fewer surface reconstruction failures and visibly better anatomical fits. At the group level, this improvement is reflected in stronger correlations between cortical thickness IDPs, indicating greater internal consistency of measurements across regions. In addition, the higher variance observed across IDPs when T2-FLAIR is used suggests that subject-specific differences are better preserved, rather than being overly constrained. Taken together, these findings are consistent with the interpretation that T1-only processing produces more conservative surface estimates, potentially driven by stronger reliance on internal priors within FreeSurfer. While increased variance in isolation could be concerning, its presence alongside improved surface quality, reduced failure rates, and stronger inter-regional correlations supports the conclusion that T2-FLAIR leads to more accurate and biologically informative estimates.

The most biased IDPs were the cortical thickness IDPs, with cortical volume and contrast also showing large mean and scale differences in many IDPs. We confirmed that the results seen in the wider UKB dataset were generalisable, with a similar distribution of Cohen’s d and *V_r_* being observed in the within-participant group and the wider sample.

While area IDPs were less affected generally, a small subset of cortical area IDPs, particularly in regions such as the parahippocampal, entorhinal, lingual, and cuneus cortices, showed biases comparable to those observed for thickness and volume measures. These regions are known to be challenging for surface reconstruction due to reduced grey matter contrast and complex local geometry. In the absence of T2-FLAIR, this can lead not only to subtle biases in pial surface placement but also to outright segmentation failures in which the surface extends into surrounding non-cortical tissue (Figures 3 & 4). Such failures can introduce a systematic outward displacement of the pial surface, resulting in overestimation of derived measures, including cortical area. The fact that these effects are concentrated in anatomically similar regions may explain why a subset of area IDPs exhibit consistent bias when T2-FLAIR is not used.

### 4.2 Method performance

#### 4.2.1 Batch effect removal

All methods that directly addressed the FST2 batch effect reduced the mean shift bias, as quantified by IDP-wise Cohen’s d, to well below a small effect size (0.2), with all methods except method 5 reducing it below a very small effect size (0.01) (Sawilowsky, 2009). Linear regression (method 2) achieved the most complete removal of the mean shift. This is expected, as OLS removes batch-specific means deterministically, whereas ComBat applies empirical Bayes shrinkage across features, resulting in a small residual mean difference. Consequently, while ComBat reduces mean differences substantially, Cohen’s d is not expected to be exactly zero. Importantly, stronger removal of an additive mean effect does not necessarily imply better harmonisation, particularly in settings with small or imbalanced sample sizes, where variance differences and overall distributional alignment are also critical.

These summaries should be interpreted as measures of alignment with the observed batch contrast rather than proof of an external ground truth, since the correction terms are estimated from the same data used to define that contrast. This circularity is most obvious for regression-based correction and for ComBat settings with relatively broad priors (e.g., Method 4), where the observed bias is removed directly in the fitted model. A stronger reduction in an additive mean effect or in variance ratios, therefore, does not by itself imply better harmonisation; assessment also requires checking distributional agreement, for example, with IDP-wise KS tests, and preservation of correlations with non-imaging variables.

The advantage of ComBat over regression for batch effect removal is then more apparent when examining variance differences between batches or using metrics that jointly assess location and scale. The batch processed without T2-FLAIR showed substantially reduced variance relative to the batch processed with T2-FLAIR, particularly in cortical thickness IDPs. Methods 1–3, which did not include scaling correction, failed to substantially improve this imbalance, with variance ratios remaining largely unchanged across IDPs. In contrast, methods 4–6 substantially improved variance agreement between batches, with mean variance ratios approaching 1 across all IDPs. However, the relative performance of the ComBat approaches differed across IDP classes. Method 4 performed best for cortical area, intensity, and grey-white matter contrast IDPs, whereas methods 5 and 6 performed best for cortical thickness IDPs, where the batch effect was strongest.

These differences likely reflect the interaction between feature heterogeneity and the empirical Bayes prior structure used by ComBat. In Method 4, all IDPs contribute to a shared global prior, producing broader priors and weaker shrinkage of the feature-wise estimates. In contrast, methods 5 and 6 estimate priors separately within IDP classes, resulting in stronger class-specific shrinkage. This appears advantageous when batch effects are relatively homogeneous within a feature class, as observed for cortical thickness IDPs, but less effective when only a subset of features within a class exhibits strong batch effects, as seen for cortical area IDPs.

Consistent with these findings, methods 4–6 substantially reduced distributional differences between batches in the two-sample Kolmogorov-Smirnov tests, whereas methods 2 and 3, which corrected only additive effects, left a substantial number of IDPs significantly different between batches. These results indicate that removal of mean shifts alone is insufficient to achieve full harmonisation when substantial variance differences are present.

No single harmonisation approach optimised all evaluation metrics simultaneously. Linear regression (Method 2) removed additive bias most completely but did not address variance differences. Among ComBat approaches, methods 5 and 6 performed best for cortical thickness IDPs, where the FST2 effect was strongest, but underperformed on the area IDPs compared to method 4. When comparing the results on the reduced sample, method 6 showed performance more consistent with that observed when the full sample was used.

#### 4.2.2 Prior distributions and IDP-wise likelihoods

Viewing the empirical priors for methods 4 and 5 provides an insight into why method 4 may seemingly outperform method 5 in some classes of IDPs but underperforms on the scaling correction for the worst affected classes. Figure 10 suggests that the influence of the empirical Bayes prior depends strongly on how well the prior distribution aligns with the feature-wise likelihood, as well as the width of the priors. In Method 4, the globally pooled prior is comparatively broad relative to the likelihood functions, meaning that the posterior estimates are dominated primarily by the feature-wise estimates and therefore are relatively less affected by the priors. By contrast, the class-specific priors in Method 5 are generally more concentrated and structured around the characteristic behaviour of their respective IDP classes, allowing them to exert greater influence on the posterior estimate. This produces stronger class-dependent shrinkage, which may improve correction in feature classes with systematic batch effects, such as thickness measures, but may also lead to under- or over-correction when the class-specific priors are estimated from smaller or more heterogeneous subsets of IDPs.

For thickness, Method 5 is the most favourable because its class-specific prior is better positioned relative to the likelihood than the pooled prior, so it supports a correction that better matches the underlying batch effect in this subgroup. In contrast, the pooled prior is already close to the likelihood and performs best overall, suggesting that the extra class-specific structure in Method 5 does not provide additional benefit and may instead introduce unnecessary constraints. Overall, the figure indicates that class-wise priors are advantageous only when they are well matched to the batch-effect structure of a given IDP class; otherwise, the broader pooled prior of Method 4 can be equally effective or even preferable.

This is seen in the cortical area IDPs, where only a small proportion of IDPs show strong batch effects (as shown by Figure 3), with the others having minimal differences between processing with and without T2-FLAIR. Here, the class-specific ComBat approach relatively undercorrects the batch effect on the mean and variance in these IDPs as seen in Figures 7 and 8. Consequently, the differences between Methods 4 and 5 appear to arise less from improved stability through global pooling and more from the differing degree to which the empirical priors constrain the feature-wise likelihood estimates. Generally, ComBat applied to IDPs that are of the same class should perform better, as the batch effect is more likely to be relatively homogeneous within a class compared to across classes. This was true for the cortical thickness IDPs, where methods 5 and 6 outperformed the global approach of method 4, but not for the area IDPs, where the batch effect was not homogeneous across the brain but instead only found in a small subgroup of regions. Importantly, across all examples shown, the likelihood functions are substantially narrower than either empirical prior. This indicates that the feature-wise estimates themselves contain strong information regarding the scaling parameter, limiting the degree to which the empirical Bayes prior can influence the posterior estimate.

#### 4.2.3 Correlations with nIDPs

Directly addressing the FST2 effect using methods 2–6 preserved the vast majority of correlations between IDPs and nIDPs, slightly strengthening them on average across all IDPs. However, some correlations between nIDPs and cortical thickness measures were reduced after accounting for the FST2 effect. As a result, some of the observed associations prior to harmonisation, particularly weaker or less biologically plausible ones, are likely to reflect artefactual relationships introduced by biased IDP estimates. The reduction in correlations following correction is therefore consistent with the removal of such bias. The correlations of IDPs and nIDPs that showed the largest reduction in strength were those of cortical thickness measures with caffeine consumption on the day of the scan and an ICD-10-coded variable describing unspecified disturbances of taste or smell. Further analysis, described in the supplementary materials, showed that the variable describing missing T2-FLAIR in FreeSurfer was significantly correlated with both of these nIDPs. As the main reason for T2-FLAIR not being included in FreeSurfer processing was due to scan interruption, it is unlikely that these correlation strengths were driven by real biology. Conversely, correlations between volumetric measures and bone and mineral density were strengthened after accounting for FST2, consistent with recovery of known biological relationships between brain size and skull properties.

When comparing the ComBat approaches to that of standard deconfounding using linear regression (method 2), we found that ComBat with the scaling term omitted showed strong agreement with standard regression, with neither approach being clearly favourable. When comparing method 3 to method 5, the mean difference in correlations was close to zero, but the direction of the correlations with respect to the mean showed a positive gradient in the y-axis, suggesting that the inclusion of the scaling term strengthened IDP-nIDP correlations, presumably by removing the thickness (FST2) bias more effectively.

Comparing methods 4–6, which all included the scaling term, showed similar correlation strengths in both the full and equal batch-size samples. The agreement between methods 5 and 6 was strongest in the full sample, as expected given the dominance of the much larger FST2=1 batch in the standardisation step, but remained high in the 1:1 sample. This suggests that the different full ComBat implementations have only limited impact on downstream IDP–nIDP correlations, even though they differ more clearly in their direct correction of batch mean and variance.

#### 4.2.4 Practical applications

From a practical perspective, our findings suggest that the optimal harmonisation strategy depends on both dataset composition and the downstream application. Where a biologically preferable or technically higher-quality batch can be identified, the reference-batch ComBat approach (Method 6) provides the most stable overall harmonisation across IDP classes and sample-size conditions. It also showed the most consistent performance when the sample sizes were set to be equal. However, this advantage is context-dependent: the suitability of a reference-batch approach relies on the presence of a well-defined and sufficiently large reference group, which in this case is provided by the substantially larger T2-FLAIR-available cohort.

For analyses focused specifically on cortical thickness measures, class-wise ComBat (Methods 5 and 6) showed slightly improved performance in correcting the strongest FST2-related effects. In contrast, simple regression-based correction (Method 2) effectively removed additive mean shifts but did not address multiplicative differences in variance between batches. As such, while Method 6 performs best for the UK Biobank FST2 scenario, its applicability may be limited in settings where no clear reference batch exists or where batch sizes are more balanced. In practice, UK Biobank can serve as an effective reference batch for harmonisation due to its large sample size, standardised acquisition, and high data quality. However, this approach assumes that the reference batch provides an accurate and unbiased estimate of the underlying data distribution. If this assumption is violated, harmonisation may propagate systematic biases or distort smaller or more heterogeneous datasets. Furthermore, this strategy is less suitable in scenarios where no clear high-quality reference batch exists, limiting it to scenarios where a biologically preferable or technically higher-quality (and/or significantly larger) batch can be identified. We have shown that this method still performs well when batch sizes are balanced (as shown in Figures 13 - 15), outperforming the other ComBat approaches quite considerably in this case by all metrics.

### 4.3 ComBat performance across sample size

A central finding of this work is the differing sample size requirements for reliably estimating additive and multiplicative batch effects. While additive effects converge rapidly and can be reliably estimated with relatively small sample sizes, multiplicative effects exhibit substantially greater instability and require considerably larger samples, particularly when variance differences between batches are large. This has important implications for real-world applications, especially in machine learning settings, as datasets that are large enough to correct mean shifts may still yield unreliable variance corrections, potentially introducing instability or bias into downstream models.

The average additive term at each sample size used in ComBat asymptotes quickly, consistent with previous literature, requiring a sample size of around 30–50 to reach a stable estimate of the additive batch effect (Fortin et al., 2017; Orlhac et al., 2022). While the estimated correction for the effect on the mean asymptotes rapidly in average value across repeats with sample size, the variability in this estimate across different random subsamples at a given batch size decreases more gradually, indicating that larger sample sizes are required for consistent estimation across repeated draws. Nevertheless, our observations remain consistent with previous studies.

The multiplicative (scaling) correction exhibited substantially greater sensitivity to batch size than the additive correction, with considerably larger batch sizes (>200 samples) required to obtain stable estimates. We further observed that this sensitivity is exacerbated when there are large differences in variance between batches. In all cases, the number of samples required for the variability of the scaling estimates to stabilise was substantially higher than the sample sizes reported as sufficient for achieving “successful” harmonisation in previous studies (Da-Ano et al., 2020; Fortin et al., 2017; Orlhac et al., 2022). This discrepancy is not unexpected: the empirical Bayes estimator for the scaling correction (see Johnson et al. (2007)) includes a *n_g_/*2 term in the denominator, implying that convergence toward a stable estimate occurs only gradually, approximately at a rate of *O*(1*/n*), where the rate of convergence is dependent on the batch size. Consequently, while reliable estimation and removal of additive batch effects may be achievable at relatively modest sample sizes, this does not necessarily extend to multiplicative effects, particularly when batch variances differ substantially. Researchers should therefore exercise caution when applying ComBat’s scaling correction and explicitly compare the variance of each batch against each other before and after applying ComBat.

These findings may have important implications for machine learning applications in neuroimaging, particularly in settings involving train/test splits, external validation cohorts, or federated analyses. In such scenarios, harmonisation parameters are often estimated on one dataset and applied to another, implicitly assuming that the estimated batch corrections generalise reliably across samples. While our results suggest that additive ComBat corrections are relatively stable even at modest sample sizes, the multiplicative variance corrections were substantially more variable, particularly in small, imbalanced, or strongly differing batches. This suggests that variance harmonisation may be less reliable in low-sample regimes and could potentially introduce instability or overfitting when applied in predictive modelling pipelines.

In this study we only examined bias arising from differences in cross-sectional FreeSurfer IDPs; as such, our results may not generalise to longitudinal FreeSurfer or other surface reconstruction pipelines that may use different models, something which could provide an interesting exploration in the future. Additionally, the missing T2-FLAIR constituted a relatively clean and well-defined batch effect with a known ground truth, one which we could test using within-participant “repeated” data. Real-world batch effects are often more complex, overlapping, and partially confounded with scanner, protocol, or population differences, which may limit the generalisability of the harmonisation findings. Additionally, we only focused on one modality here; as such, while some aspects of the findings would likely be generalisable to other modalities, this would require further study.

## 5 Conclusion

Including T2-FLAIR when running FreeSurfer helps to preserve real inter-subject variability, improves the correlations of thickness IDPs across different brain regions and reduces the rate of failures in estimating the pial surface by providing contrast information for regions where contrast differences may be ambiguous in the T1-weighted image. In the case of UK Biobank data, it is valuable to apply a harmonisation approach such as ComBat to IDPs affected by whether the T2-FLAIR was available in order to correct those IDPs for both shift and scaling biases.

Careful analysis of the nature of the batch effect is needed to ensure the correct approach is used. We found that when there is little to no scaling/variance difference between batches due to batch effects, using linear regression to remove batch and covariate effects is sufficient for deconfounding and harmonisation. When there is a scaling difference between batches, ComBat is beneficial, showing improved variance agreement between batches and improved results of the two-sample Kolmogorov-Smirnov test while preserving correlations with nIDPs.

Overall, across metrics and sample sizes, the reference-batch ComBat approach (Method 6) performed best in correcting both additive and multiplicative batch effects while preserving biological signal. This was most evident in IDPs with strong batch effects on the variance, particularly cortical thickness measures. However, when batch effects were heterogeneous across an IDP class, Method 4, which used global pooling, performed comparatively better, with this likely being due to the likelihood dominating the posterior estimate of the correction in these cases. While linear regression was sufficient for correcting mean shifts, it failed to address scaling differences between batches. In contrast, ComBat approaches provided a more complete harmonisation framework, correcting scaling and distributional differences while still preserving correlations with non-imaging variables.

A central finding of this work is that additive and multiplicative batch corrections behave differently with respect to sample size. While additive effects can be estimated reliably with relatively modest samples, stable estimation of multiplicative variance corrections requires substantially larger datasets, particularly when batch differences are large or imbalanced. These findings suggest that successful removal of mean differences alone should not be interpreted as evidence of complete harmonisation and highlight the need for greater caution when applying variance correction in smaller neuroimaging datasets.

## Acknowledgments

This research has been conducted in part using the UK Biobank Resource under Application Number 8107. We are grateful to UK Biobank for making the data available and to all UK Biobank study participants, who generously donated their time to make this resource possible. Analysis was carried out on the clusters at the Oxford Biomedical Research Computing (BMRC) facility and FMRIB (part of the Oxford Centre for Integrative Neuroimaging). BMRC is a joint development between the Wellcome Centre for Human Genetics and the Big Data Institute, supported by Health Data Research UK and the NIHR Oxford Biomedical Research Centre.

## Data and code availability statement

The UK Biobank data used in this study are available to approved researchers through the UK Biobank Access Management System, subject to UK Biobank’s standard access procedures. This study was conducted under UK Biobank application number 8107. Derived data that could compromise participant privacy are not publicly shared. Code implementing the ComBat approaches and analyses described in this manuscript is available through the supplementary materials.

## Authors’ Contributions

**Conceptualisation:** Jacob Turnbull, Stephen Smith, Ludovica Griffanti

**Data curation:** Jacob Turnbull, Stephen Smith, Ludovica Griffanti

**Formal analysis:** Jacob Turnbull

**Funding acquisition:** Stephen Smith, Ludovica Griffanti

**Methodology:** Jacob Turnbull, Stephen Smith, Ludovica Griffanti

**Project administration:** Jacob Turnbull, Stephen Smith, Ludovica Griffanti

**Software:** Jacob Turnbull, Fidel Alfaro-Almagro and Stephen Smith

**Supervision:** Gaurav Bhalerao, Stephen Smith, Ludovica Griffanti

**Visualisation:** Jacob Turnbull, Rach Dawson, Frederik Lange

**Writing – original draft:** Jacob Turnbull

**Writing – review & editing:** Jacob Turnbull, Gaurav Bhalerao, Rach Dawson, Frederik Lange, Fidel Alfaro-Almagro, Stephen Smith, Ludovica Griffanti

## Supplementary materials

The modified version of ComBat used in the analysis of this project is publicly available on GitHub in both Python and MATLAB. We additionally provide the code required to create most of the figures from this work: (https://github.com/Jake-Turnbull/ComBat_FST2_paper).

## References

Alfaro-Almagro, F. (2021, August). Brain imaging in UK Biobank [PhD thesis]. (Cit. on pp. 3, 4).

Alfaro-Almagro, F., Jenkinson, M., Bangerter, N. K., Andersson, J. L. R., Griffanti, L., Douaud, G., Sotiropoulos, S. N., Jbabdi, S., Hernandez-Fernandez, M., Vallee, E., Vidaurre, D., Webster, M., McCarthy, P., Rorden, C., Daducci, A., Alexander, D. C., Zhang, H., Dragonu, I., Matthews, P. M., . . . Smith, S. M. (2018). Image processing and Quality Control for the first 10,000 brain imaging datasets from UK Biobank. NeuroImage, 166, 400–424 (cit. on p. 3).

Alfaro-Almagro, F., McCarthy, P., Afyouni, S., Andersson, J. L. R., Bastiani, M., Miller, K. L., Nichols, T. E., & Smith, S. M. (2021). Confound modelling in UK Biobank brain imaging. NeuroImage, 224, 117002 (cit. on pp. 6, 11, 33).

Bayer, J. M. M., Thompson, P. M., Ching, C. R. K., Liu, M., Chen, A., Panzenhagen, A. C., Jahanshad, N., Marquand, A., Schmaal, L., & Sämann, P. G. (2022). Site effects how-to and when: An overview of retrospective techniques to accommodate site effects in multi-site neuroimaging analyses. Frontiers in Neurology, 13 (cit. on p. 3).

Carré, A., Battistella, E., Niyoteka, S., Sun, R., Deutsch, E., & Robert, C. (2022). AutoComBat: A generic method for harmonizing MRI-based radiomic features. Scientific Reports, 12 (1), 12762 (cit. on p. 7).

Cetin Karayumak, S., Bouix, S., Ning, L., James, A., Crow, T., Shenton, M., Kubicki, M., & Rathi, Y. (2019). Retrospective harmonization of multi-site diffusion MRI data acquired with different acquisition parameters. NeuroImage, 184, 180–200 (cit. on p. 2).

Corey, D., Dunlap, W., & Burke, M. (1998). Averaging Correlations: Expected Values and Bias in Combined Pearson rs and Fisher’s z Transformations. Journal of General Psychology - J GEN PSYCHOL, 125, 245–261 (cit. on p. 11).

Da-Ano, R., Masson, I., Lucia, F., Doré, M., Robin, P., Alfieri, J., Rousseau, C., Mervoyer, A., Reinhold, C., Castelli, J., De Crevoisier, R., Rameé, J. F., Pradier, O., Schick, U., Visvikis, D., & Hatt, M. (2020). Performance comparison of modified ComBat for harmonization of radiomic features for multicenter studies. Scientific Reports, 10 (1), 10248 (cit. on p. 37).

Dewey, B. E., He, Y., Liu, Y., Zuo, L., & Prince, J. L. (2022, January). Chapter 11 - Medical image harmonization through synthesis. In N. Burgos & D. Svoboda (Eds.), Biomedical Image Synthesis and Simulation (pp. 217–232). Academic Press. (Cit. on p. 10).

Elliott, L. T., Sharp, K., Alfaro-Almagro, F., Shi, S., Miller, K., Douaud, G., Marchini, J., & Smith, S. (2018, June). Genome-wide association studies of brain structure and function in the UK Biobank [Pages: 178806 Section: New Results]. (Cit. on p. 3).

Ferreira-Atuesta, C., Binder, S., Piendel, L. A., Emerson, J. M., Andrew-Jaja, N., Sutkowski, J., & Hedden, T. (2022). Comparison of FreeSurfer inputs on estimates and reliability of volume and thickness measures. Alzheimer’s & Dementia, 18 (S5), e066079 (cit. on p. 3).

Fischl, B. (2012). FreeSurfer. NeuroImage, 62 (2), 774–781 (cit. on pp. 3, 33).

Fischl, B., Salat, D. H., Busa, E., Albert, M., Dieterich, M., Haselgrove, C., van der Kouwe, A., Killiany, R., Kennedy, D., Klaveness, S., Montillo, A., Makris, N., Rosen, B., & Dale, A. M. (2002). Whole brain segmentation: Automated labeling of neuroanatomical structures in the human brain. Neuron, 33 (3), 341–355 (cit. on p. 5).

Fortin, J.-P. (2021, July). Jfortin1/ComBatHarmonization [original-date: 2017-03-23T14:15:19Z]. (Cit. on p. 7).

Fortin, J.-P., Cullen, N., Sheline, Y. I., Taylor, W. D., Aselcioglu, I., Cook, P. A., Adams, P., Cooper, C., Fava, M., McGrath, P. J., McInnis, M., Phillips, M. L., Trivedi, M. H., Weissman, M. M., & Shinohara, R. T. (2018). Harmonization of cortical thickness measurements across scanners and sites. NeuroImage, 167, 104–120 (cit. on pp. 4, 7).

Fortin, J.-P., Parker, D., Tunç, B., Watanabe, T., Elliott, M. A., Ruparel, K., Roalf, D. R., Satterthwaite, T. D., Gur, R. C., Gur, R. E., Schultz, R. T., Verma, R., & Shinohara, R. T. (2017). Harmonization of multi-site diffusion tensor imaging data. NeuroImage, 161, 149–170 (cit. on pp. 7, 12, 32, 37).

Giavarina, D. (2015). Understanding Bland Altman analysis. Biochemia Medica, 25 (2), 141–151 (cit. on p. 11).

Habes, M., Pomponio, R., Shou, H., Doshi, J., Mamourian, E., Erus, G., Nasrallah, I., Launer, L. J., Rashid, T., Bilgel, M., Fan, Y., Toledo, J. B., Yaffe, K., Sotiras, A., Srinivasan, D., Espeland, M., Masters, C., Maruff, P., Fripp, J., . . . iSTAGING consortium, the Preclinical AD consortium, the ADNI, and the CARDIA studies. (2021). The Brain Chart of Aging: Machine-learning analytics reveals links between brain aging, white matter disease, amyloid burden, and cognition in the iSTAGING consortium of 10,216 harmonized MR scans. Alzheimer’s & Dementia: The Journal of the Alzheimer’s Association, 17 (1), 89–102 (cit. on p. 2).

Iglesias, J. E., Insausti, R., Lerma-Usabiaga, G., Bocchetta, M., Van Leemput, K., Greve, D. N., van der Kouwe, A., Alzheimer’s Disease Neuroimaging Initiative, Fischl, B., Caballero-Gaudes, C., & Paz-Alonso, P. M. (2018). A probabilistic atlas of the human thalamic nuclei combining ex vivo MRI and histology. NeuroImage, 183, 314–326 (cit. on p. 3).

Jodoin, P.-M., Edde, M., Girard, G., Dumais, F., Theaud, G., Dumont, M., Houde, J.-C., David, Y., & Descoteaux, M. (2025). Challenges and best practices when using ComBAT to harmonize diffusion MRI data. Scientific Reports, 15 (1), 41508 (cit. on p. 32).

Johnson, W. E., Li, C., & Rabinovic, A. (2007). Adjusting batch effects in microarray expression data using empirical Bayes methods. Biostatistics, 8 (1), 118–127 (cit. on pp. 7, 8, 37).

Klapwijk, E. T., van de Kamp, F., van der Meulen, M., Peters, S., & Wierenga, L. M. (2019). Qoala-T: A supervised-learning tool for quality control of FreeSurfer segmented MRI data. NeuroImage, 189, 116–129 (cit. on pp. 4, 33).

Lindroth, H., Nair, V. A., Stanfield, C., Casey, C., Mohanty, R., Wayer, D., Rowley, P., Brown, R., Prabhakaran, V., & Sanders, R. D. (2019). Examining the identification of age-related atrophy between T1 and T1 + T2-FLAIR cortical thickness measurements. Scientific Reports, 9 (1), 11288 (cit. on pp. 3, 6, 33).

Lu, Y.-C., Zuo, L., Chou, Y.-Y., Dewey, B. E., Remedios, S., Shinohara, R. T., Steele, S. U., Nair, G., Reich, D. S., Prince, J. L., & Pham, D. L. (2025). An Evaluation of Image-Based and Statistical Techniques for Harmonizing Brain Volume Measurements. Imaging Neuroscience (cit. on p. 10).

Marzi, C., Giannelli, M., Barucci, A., Tessa, C., Mascalchi, M., & Diciotti, S. (2024). Efficacy of MRI data harmonization in the age of machine learning: A multicenter study across 36 datasets. Scientific Data, 11 (1), 115 (cit. on pp. 4, 11).

Miller, K. L., Alfaro-Almagro, F., Bangerter, N. K., Thomas, D. L., Yacoub, E., Xu, J., Bartsch, A. J., Jbabdi, S., Sotiropoulos, S. N., Andersson, J. L. R., Griffanti, L., Douaud, G., Okell, T. W., Weale, P., Dragonu, I., Garratt, S., Hudson, S., Collins, R., Jenkinson, M., . . . Smith, S. M. (2016). Multimodal population brain imaging in the UK Biobank prospective epidemiological study. Nature Neuroscience, 19 (11), 1523–1536 (cit. on p. 2).

Newman-Norlund, R. D., Newman-Norlund, S. E., Sayers, S., Nemati, S., Riccardi, N., Rorden, C., & Fridriksson, J. (2021). The Aging Brain Cohort (ABC) repository: The University of South Carolina’s multimodal lifespan database for studying the relationship between the brain, cognition, genetics and behavior in healthy aging. Neuroimage: Reports, 1 (1), 100008 (cit. on p. 2).

Orlhac, F., Eertink, J. J., Cottereau, A.-S., Zijlstra, J. M., Thieblemont, C., Meignan, M., Boellaard, R., & Buvat, I. (2022). A Guide to ComBat Harmonization of Imaging Biomarkers in Multicenter Studies. Journal of Nuclear Medicine, 63 (2), 172–179 (cit. on pp. 4, 7, 12, 32, 37).

Parekh, P., Vivek Bhalerao, G., Viswanath, B., Rao, N. P., Narayanaswamy, J. C., Sivakumar, P. T., Kandasamy, A., Kesavan, M., Mehta, U. M., Mukherjee, O., Purushottam, M., Mehta, B., Kandavel, T., Binukumar, B., Saini, J., Jayarajan, D., Shyamsundar, A., Moirangthem, S., Vijay Kumar, K. G., . . . Venkatasubramanian, G. (2022). Sample size requirement for achieving multisite harmonization using structural brain MRI features. NeuroImage, 264, 119768 (cit. on p. 4).

Petersen, R. C., Aisen, P. S., Beckett, L. A., Donohue, M. C., Gamst, A. C., Harvey, D. J., Jack, C. R., Jagust, W. J., Shaw, L. M., Toga, A. W., Trojanowski, J. Q., & Weiner, M. W. (2010). Alzheimer’s Disease Neuroimaging Initiative (ADNI). Neurology, 74 (3), 201–209 (cit. on p. 2).

Pomponio, R., Erus, G., Habes, M., Doshi, J., Srinivasan, D., Mamourian, E., Bashyam, V., Nasrallah, I. M., Satterthwaite, T. D., Fan, Y., Launer, L. J., Masters, C. L., Maruff, P., Zhuo, C., Völzke, H., Johnson, S. C., Fripp, J., Koutsouleris, N., Wolf, D. H., . . . Davatzikos, C. (2020). Harmonization of large MRI datasets for the analysis of brain imaging patterns throughout the lifespan. NeuroImage, 208, 116450 (cit. on p. 7).

Radua, J., Vieta, E., Shinohara, R., Kochunov, P., Quidé, Y., Green, M. J., Weickert, C. S., Weickert, T., Bruggemann, J., Kircher, T., Nenadić, I., Cairns, M. J., Seal, M., Schall, U., Henskens, F., Fullerton, J. M., Mowry, B., Pantelis, C., Lenroot, R., . . . Pineda-Zapata, J. (2020). Increased power by harmonizing structural MRI site differences with the ComBat batch adjustment method in ENIGMA. NeuroImage, 218, 116956 (cit. on p. 8).

Riffenburgh, R. H., & Gillen, D. L. (2020a, January). 13 - Tests on variability and distributions. In R. H. Riffenburgh & D. L. Gillen (Eds.), Statistics in Medicine (Fourth *Edition)* (pp. 311–336). Academic Press. (Cit. on p. 10).

Riffenburgh, R. H., & Gillen, D. L. (2020b, January). 27 - Techniques to Aid Analysis. In R. H. Riffenburgh & D. L. Gillen (Eds.), Statistics in Medicine (Fourth *Edition)* (pp. 631–649). Academic Press. (Cit. on p. 5).

Sawilowsky, S. S. (2009). New Effect Size Rules of Thumb. Journal of Modern Applied Statistical Methods, 8 (2), 597–599 (cit. on pp. 25, 34).

Smith, S., Alfaro-Almagro, F., & Miller, K. (2025, May). UK Biobank Brain Imaging Documentation. (Cit. on p. 17).

Smith, S. M., & Nichols, T. E. (2018). Statistical Challenges in “Big Data” Human Neuroimaging. Neuron, 97 (2), 263–268 (cit. on p. 2).

Stein, C. K., Qu, P., Epstein, J., Buros, A., Rosenthal, A., Crowley, J., Morgan, G., & Barlogie, B. (2015). Removing batch effects from purified plasma cell gene expression microarrays with modified ComBat. BMC Bioinformatics, 16 (1), 63 (cit. on p. 9).

Stroganov, O., Fedarovich, A., Wong, E., Skovpen, Y., Pakhomova, E., Grishagin, I., Fedarovich, D., Khasanova, T., Merberg, D., Szalma, S., & Bryant, J. (2022). Mapping of UK Biobank clinical codes: Challenges and possible solutions. PLOS ONE, 17 (12), e0275816 (cit. on p. 24).

Sudlow, C., Gallacher, J., Allen, N., Beral, V., Burton, P., Danesh, J., Downey, P., Elliott, P., Green, J., Landray, M., Liu, B., Matthews, P., Ong, G., Pell, J., Silman, A., Young, A., Sprosen, T., Peakman, T., & Collins, R. (2015). UK Biobank: An Open Access Resource for Identifying the Causes of a Wide Range of Complex Diseases of Middle and Old Age. PLoS Medicine, 12 (3), e1001779 (cit. on pp. 2, 24).

Thompson, P. M., Jahanshad, N., Ching, C. R. K., Salminen, L. E., Thomopoulos, S. I., Bright, J., Baune, B. T., Bertolín, S., Bralten, J., Bruin, W. B., Bülow, R., Chen, J., Chye, Y., Dannlowski, U., de Kovel, C. G. F., Donohoe, G., Eyler, L. T., Faraone, S. V., Favre, P., . . . Zelman, V. (2020). ENIGMA and global neuroscience: A decade of large-scale studies of the brain in health and disease across more than 40 countries. Translational Psychiatry, 10 (1), 100 (cit. on p. 2).

Van Essen, D. C., Ugurbil, K., Auerbach, E., Barch, D., Behrens, T. E. J., Bucholz, R., Chang, A., Chen, L., Corbetta, M., Curtiss, S. W., Della Penna, S., Feinberg, D., Glasser, M. F., Harel, N., Heath, A. C., Larson-Prior, L., Marcus, D., Michalareas, G., Moeller, S., . . . WU-Minn HCP Consortium. (2012). The Human Connectome Project: A data acquisition perspective. NeuroImage, 62 (4), 2222–2231 (cit. on p. 2).

